# Multiplexed characterization of rationally designed promoter architectures deconstructs combinatorial logic for IPTG-inducible systems

**DOI:** 10.1101/2020.01.31.928689

**Authors:** Timothy C. Yu, Winnie L. Liu, Marcia Brinck, Jessica E. Davis, Jeremy Shek, Grace Bower, Tal Einav, Kimberly D. Insigne, Rob Phillips, Sriram Kosuri, Guillaume Urtecho

**Author notes:** The authors wish it to be known that, in their opinion, the first two authors should be regarded as joint First Authors.

## Abstract

A crucial step towards engineering biological systems is the ability to precisely tune the genetic response to environmental stimuli. In the case of *Escherichia coli* inducible promoters, our incomplete understanding of the relationship between sequence composition and gene expression hinders our ability to predictably control transcriptional responses. Here, we profile the expression dynamics of 8,269 rationally designed IPTG-inducible promoters that collectively explore the individual and combinatorial effects of RNA polymerase and LacI repressor binding site strengths. Using these data, we fit a statistical mechanics model that accurately models gene expression and reveals properties of theoretically optimal inducible promoters. Furthermore, we characterize three novel promoter architectures and show that repositioning binding sites within promoters influences the types of combinatorial effects observed between promoter elements. In total, this approach enables us to deconstruct relationships between inducible promoter elements and discover practical insights for engineering inducible promoters with desirable characteristics.

## Introduction

Inducible promoters are key regulators of cellular responses to external stimuli and popular engineering targets for applications in metabolic flux optimization and biosensing^1–3^. For example, inducible systems have been designed to function as controlled cell factories for chemical biosynthesis as well as non-invasive diagnostics for gut inflammation^4,5^. However, these applications generally rely on synthetic inducible promoters that can elicit precisely programmable responses, a quality that is not exhibited by native promoter systems. As a result, there is a demand for new strategies to engineer customizable inducible promoters with desirable characteristics, such as minimal expression in the uninduced state (minimal leakiness) and maximal difference between the induced and uninduced states (maximal fold-change). More broadly, the design and analysis of synthetic inducible promoter function provides insight on the biophysical processes driving gene regulation.

A variety of compelling approaches have been implemented to engineer inducible promoters, however, these strategies have their shortcomings. Previous studies have had great success implementing biophysical models to tune the relative behaviors of regulatory elements and explain promoter expression, but these do not tell us how the repositioning of binding sites influences expression^6–10^. Directed evolution is a promising strategy that leverages stepwise random mutagenesis and selection to identify favorable promoters, but is generally limited to optimizing within local, evolutionarily accessible sequence space^11,12^. While this ‘black box’ approach can produce variants with the desired phenotype, it often requires iterative rounds of library screenings^12^ and does not inform our ability to logically construct promoters. Lastly, rational design is a useful approach based on the application of pre-existing mechanistic knowledge of inducible systems to generate novel variants^13,14^. Although there is great potential in rationally designed promoters for achieving specific applications, a caveat is that this approach requires a fundamental understanding of how to engineer these systems.

Inducible promoters consist of cis-regulatory elements that work in concert with multiple *trans*-acting factors to determine overall expression output15,16. As such, a critical step toward learning how to engineer these systems is to interrogate the combinatorial regulatory effects between promoter-based elements. Years of studies on the inducible *lacZYA* promoter have revealed many sequence-based factors influencing its regulation and expression. First, the binding affinities of operator sites are critical elements in determining the activity of the repressor protein, LacI^17,18^. Second, the nucleotide spacing between operator sites is vital as looping-mediated repression is dependent on repressor orientation^17,19^. Third, the positioning of the repressor sites relative to the RNAP binding sites determine a variety of repression mechanisms and transcriptional behaviors^13,14^. Fourth, the strength of the core promoter modulates RNA polymerase avidity and thus resultant gene expression^6^. However, while previous studies have characterized these modular sequence components individually, the combinatorial effects of these features on promoter induction have yet to be explored.

Inspired by previous success in studying the combinatorial logic of *E. coli* promoters^20^, we sought to address these obstacles by integrating rational design with high-throughput screening of large DNA-encoded libraries. The recent development of massively-parallel reporter assays (MPRAs) provides a framework for leveraging next-generation sequencing to measure cellular transcription levels of large numbers of DNA sequence variants. This approach enables us to measure the activity of thousands of synthetic sequences in a single, multiplexed experiment using transcriptional barcodes as a readout^20,21^. Here, we implemented a genomically-encoded MPRA system to interrogate thousands of rationally designed variants of the *lacZYA* promoter and investigate relationships between inducible promoter components across four sequence architectures. We first explore the relationship between operator spacing and repression at the *lacUV5* promoter using a variety of transcriptional repressors. Next, we designed and characterized 8,269 promoters composed of combinations of LacI repressor and RNAP binding sites, exploring combinatorial interactions between elements and establishing relationships that guide transcriptional behavior. Lastly, we isolated and further characterized promoters with various levels of fold-change and leakiness that may be useful in synthetic applications.

## Results

### Repression by transcription factors is dependent on operator spacing

The *lacZYA* promoter is a classic model for gene regulation in *E. coli*, with many studies investigating the relationship between sequence composition and induction properties. This promoter contains two LacI dimer sites positioned at the *proximal* +11 and *distal* −82 positions relative to the transcription start site (TSS)^22,23^, which flank a set of σ70 −10 and −35 elements. RNAP cooperatively binds these σ70 hexameric sequences and the relative binding affinity of these elements determines the strength of the promoter^6,8^. Conversely, the LacI operator sites repress the native *lacZYA* promoter when bound^24^. While LacI repressor bound at the *proximal* site blocks RNAP binding as well as promoter escape, binding at the *distal* site alone does not inhibit transcription and serves a more nuanced role in repression^25^. When both the *proximal* and *distal* sites are bound, LacI dimers at these sites can engage in a homotetrameric protein interaction, tethering these sites together and forming a local DNA loop^18,26,27^. This ‘repression loop’ further occludes RNAP binding, decreasing gene expression.

Studies exploring the formation of this repression loop have found that it is heavily dependent on the spacing of the LacI operator sites relative to each other^28–30^. Due to the helical nature of B-form DNA, which completes a full rotation roughly every 10.5 bp, as operator sites are placed at various distances from one another along the DNA this also changes their relative orientation along the face of the DNA helix. As a result, the ability of the *distal* site to engage in this repression loop fluctuates as it is shifted across the promoter, with repression strength correlated with helical phasing between the two operator sites^28,29^. In our effort to optimize the *lacZYA* promoter, we sought to validate the effect that operator spacing has on repression, as well as explore whether other repressors follow this same phenomenon.

To explore this, we tested the relationship between spacing and repression for six transcription factors (TFs) at the *lacUV5* promoter: LacI, AraC, GalR, GlpR, LldR, and PurR. While only LacI^27,29,31^, AraC^32,33^ and GalR^34–38^ have been experimentally shown to engage in DNA looping, there is evidence that GlpR^39^, LldR^40^, and PurR^26^ may also be capable of this mechanism. Using reported, natural binding sites for these transcription factors^41^, we designed 624 sequences assessing the ability of these sites to repress a constitutive *lacUV5* across various operator spacings. In our design, a *proximal* site for each transcription factor was centered at +12, to avoid sites overlapping the transcription start site, and a series of variants were created in which the *distal* operator site was centered at each position from −83 to −116 relative to the TSS **(Figure 1A)**. Furthermore, to quantify the effect of the individual sites, we tested variants where either the *proximal* or *distal* site was replaced with a scrambled sequence variant that maintained the GC content of each site. We grew this library in MOPS rich-defined media supplemented with 0.2% glucose, a condition for which all transcription factors should be repressive, and measured expression of all variants using a previously described MPRA^20^ **(Figure 1B)**. In brief we synthesized each variant and engineered these promoters to express uniquely barcoded GFP transcripts. Using recombination-mediated cassette exchange^42^, each barcoded variant is singly integrated into the *essQ-cspB* intergenic locus of the *E. coli* genome, positioned near the chromosomal midreplichore. We then grew the integrated libraries in rich, defined media and quantified relative barcode expression levels by performing RNA-Seq of the transcribed barcodes and normalizing transcript levels to DNA copy number as determined by DNA-Seq. Using this assay, we recovered expression measurements for 615 (98.6%) of the variants we designed, measuring an average of 70 unique barcodes per variant (**Figure S1**). These measurements exhibited a high degree of correlation between technical replicates (**Figure 1C,** *r* = 0.987, *p* < 2.2 × −10^16^).

**Figure 1.**
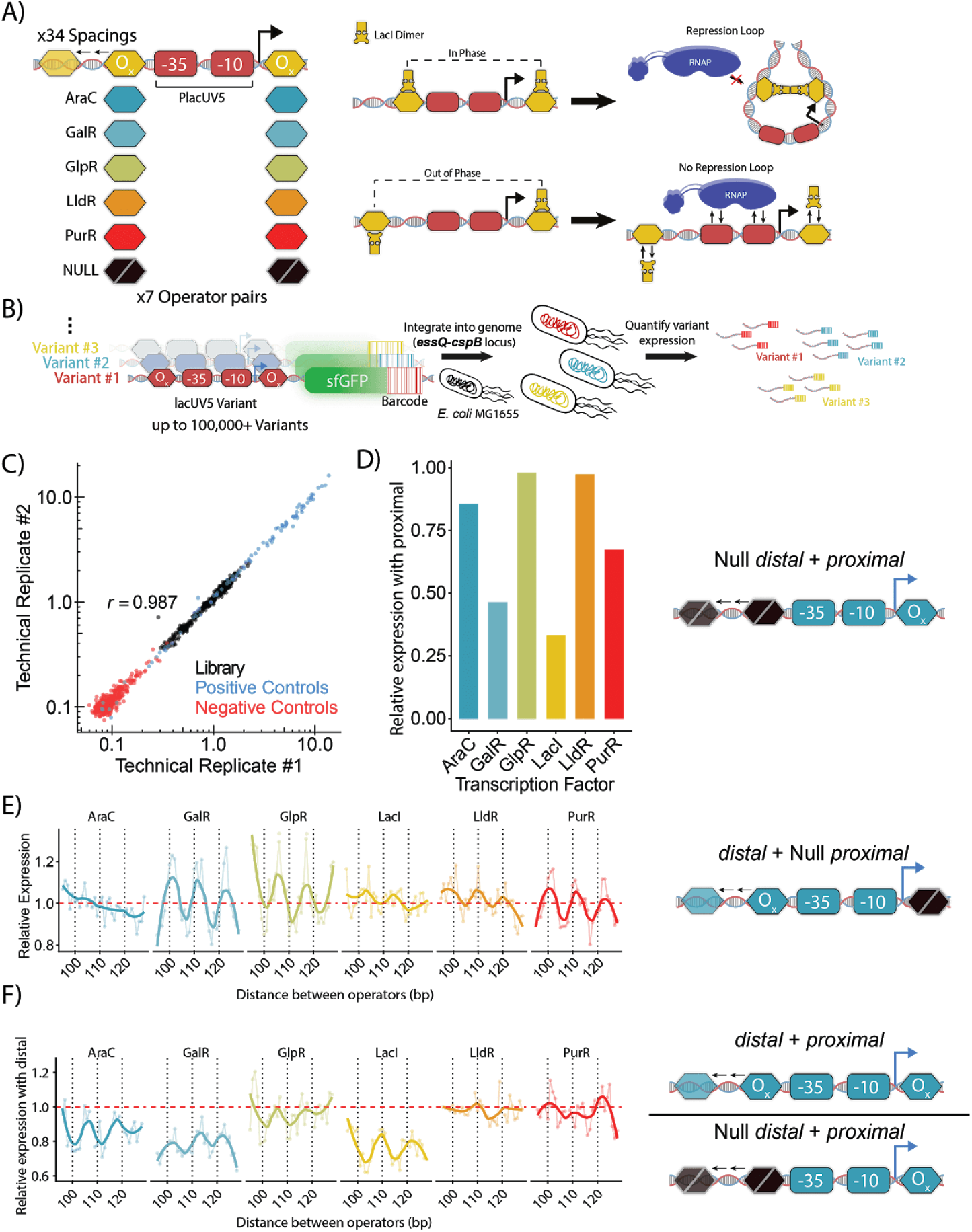
Identifying optimal spacing for repressors at *lacUV5* promoter. **A)** We designed a library of *lacUV5* variants to test repressor effects when the *distal* site was moved 32 nucleotides upstream at 1 bp increments. If repressors bind along the same face of the DNA helix, repression loop formation may occur, thereby preventing RNAP association with the promoter. **B)** MPRA for measuring promoter activity of up to hundreds of thousands of synthesized variants. Pooled promoter variants are engineered to expression uniquely barcoded sfGFP transcripts, singly integrated into the *essQ-cspB* locus of the *E. coli* genome, and characterized by quantitative RNA-Seq of the barcodes. **C)** Comparison of expression measurements between biological replicates grown in MOPS rich-defined medium supplemented with 0.2% glucose (*r* = 0.987, p < 2.2 × 10^−16^). **D)** Expression when *proximal* site is added relative to expression of *lacUV5* without repressor sites. **E)** Relative expression as each *distal* site is moved upstream in the absence of a *proximal* site relative to *lacUV5* without repressors. Thick lines denote the fit using locally weighted polynomial regression. Thin lines connect data points at sequential intervals. **F)** Relative expression as the *distal* site is moved upstream when the *proximal* site is present relative to expression of the *proximal*-only variant.

We first explored the ability of these TFs to repress the *lacUV5* promoter when placed in the *proximal* position. To evaluate this, we compared the relative difference in expression between variants with *proximal* sites to the *lacUV5* promoter with LacI sites removed (**Figure 1D**). At this position, repression varied across operators although the LldR and GlpR sites were entirely ineffective. LacI exhibited the strongest level of repression in the *proximal* position at 2.62-fold, which may be due to the strong binding affinity of the native *proximal* operator site^22^.

To gauge the performance of these repressors at each position in the *distal* site, we looked at how expression changes as a function of increased distance from the *proximal* site. While LacI^43^ and AraC^32,33^ are known to exhibit a cyclic pattern of repression as the distance between operator sites is increased, there are no direct measurements showing that GalR, GlpR, LldR, or PurR share this phenomenon. First, we looked at the effect of moving the *distal* site across 33 nucleotides in the absence of a functional *proximal* site (**Figure 1E**). We observed a uniformity of response across all repressors tested, suggesting cyclic repression is a general phenomenon of many transcription factors. Interestingly, several of these transcription factors alternated between repression and activation effects on the promoter depending on their position. Conversely, AraC binding sites gradually increased repression as they moved further upstream, with a significant inverse relationship between operator distance and expression, though the effect size is small (*p* = 2.19 × 10^−5^, ANOVA). To see whether these relationships would change when DNA looping was possible, we evaluated the effect of moving the *distal* site when the *proximal* site was also present (**Figure 1F**). To directly observe the impact of the *distal* site, we determined the expression at each *distal* position relative to expression when only the *proximal* site was present. Coupled with a *proximal* site, a majority of tested transcription factors exhibited different repression patterns as the *distal* site was moved further upstream. For AraC, GalR, and LacI the *distal* sites reduce expression more with a *proximal* site present than without (AraC: 1.18-fold, *p* = 1.83 × 10^−8^, Welch’s t-test; GalR: 1.35-fold, *p* = 2.82 × 10^−11^, Welch’s t-test; LacI: 1.37-fold, *p* = 4.65 × 10^−14^, Welch’s t-test). This enhanced repression by *distal* sites when a *proximal* site is present indicates the existence of synergistic interactions between these sites. Furthermore, repression by these *distal* sites followed a 10-11 bp periodicity as they were moved incrementally further from the *proximal* site, which may indicate the formation of DNA loops at the *lacUV5* promoter. LldR, PurR, and GlpR *distal* sites did not show show significantly enhanced ability to repress when a *proximal* site was present (*p* > 0.4 in all cases, Welch’s t-test). Instead they demonstrated similar levels of repression at all positions when a *proximal* site was present. Thus, we find that different repressor systems exhibit unique relationships between operator spacings and repression, highlighting the need to study these systems individually.

### Tuning binding site strengths alters inducible promoter behavior

We deepened our investigation into the *lacUV5* promoter to explore how the strength of RNAP and LacI binding sites contribute to its behavior. In particular, we sought to learn how these sites may be manipulated to generate *lacUV5* variants with minimal leakiness and maximal fold-change, properties that are generally desired in synthetic applications. In previous work, we have found that testing large libraries of promoters composed of various combinations of sequence elements allows us to characterize the contribution of individual sequence elements and reveal interactions between these elements^20,44^. We utilized our MPRA to uncover how tuning binding sites within the *lacUV5* promoter affects the fold-change and leakiness of this system. We designed and assayed a library of 1,600 inducible promoters, which we refer to as Pcombo, composed of all possible combinations of one of ten *proximal* LacI binding sites at +11, four −10 elements, four −35 elements, and ten *distal* LacI sites at −90 **(Figure 2A)**. To cover a wide range of expression, we selected −10 and −35 element variants previously shown to span a range of RNAP binding affinities^6,20,44^. Similarly, we chose a range of LacI binding site variants from well-characterized genomic operator sites (O_1_, O_3_, O_sym_)^10,18^, a variant of the natural O_2_ site, O_2-var_, and a series of novel LacI sites created from different combinations of the monomeric halves of each of these dimeric binding sites (**Table S2**). While O_1_ is the naturally occurring operator site reported to have the highest affinity for LacI, the synthetic O_sym_ is a symmetrized variant with even higher affinity^18,45^. Expression data for these variants was collected in both uninduced (0 mM IPTG) and fully induced conditions (1 mM IPTG). We recovered expression measurements for 1,493 variants within this library (93.3%) with an average of 9 barcodes measured per variant. We observed high expression correlation between biological replicates in both the induced and uninduced conditions (Induced: *r* = 0.945, *p* < 2.2 × 10^−16^, Uninduced: *r* = 0.955, *p* < 2.2 × 10^−16^, Welch’s t-test) (**Figure S2A**).

**Figure 2.**
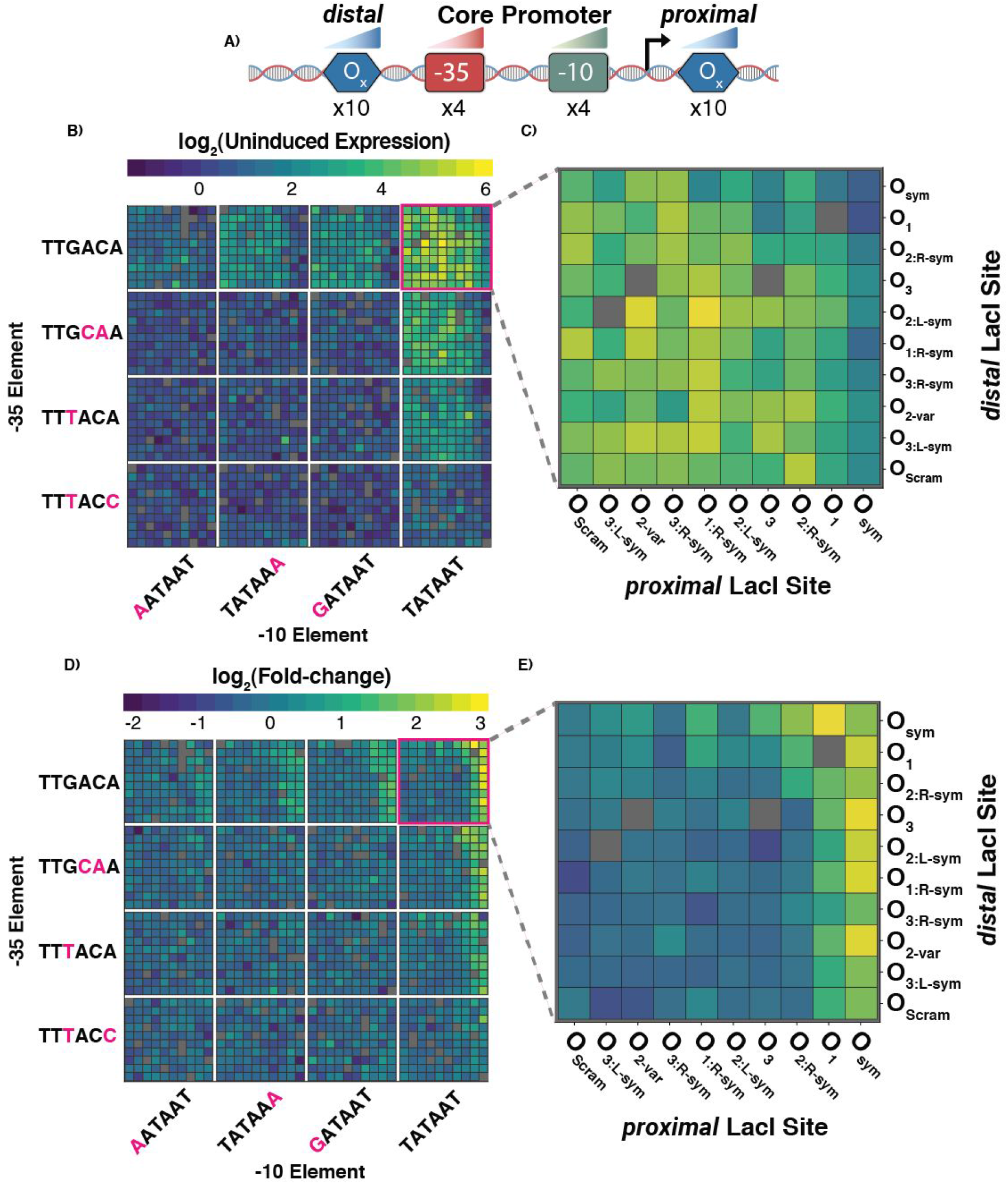
Tuning binding site strengths alters inducible promoter behavior. **A)** Pcombo library schematic consists of all combinations of one of ten *proximal* LacI binding sites, four −10 elements, four −35 elements, and ten *distal* LacI sites. **B)** Uninduced expression for all assayed Pcombo variants. Grid positions for the −10 and −35 motifs are arranged from weakest tested sequence to the consensus (−10:TATAAT and −35:TTGACA). Gray boxes indicate sequences which were not measured by the assay. **C)** Uninduced expression for assayed Pcombo variants containing a consensus core promoter. **D)** Fold-change for all assayed Pcombo variants. Fold-change is determined by the ratio of expression at 1 mM IPTG relative to 0 mM IPTG. **E)** Fold-change for all assayed Pcombo variants containing a consensus core promoter.

We first explored how the composition of sequence elements determined uninduced promoter expression, or leakiness. Library variants exhibited a 267-fold range of expression in the uninduced state overall, and even amongst variants containing the same core promoter σ70 elements expression varied by up to 96-fold (**Figure 2B**). In this uninduced state, promoters composed of the consensus −10/−35 elements exhibited the greatest leakiness, with up to 21-fold higher average expression than that of promoters composed of weaker −10/−35 elements. Effective repression generally required a strong LacI operator site, such as O_sym_ and O_1_, in the *proximal* position, especially amongst variants with consensus −10/−35 elements (**Figure 2C**).

Next, we compared the induced and uninduced expression of each *lacUV5* variant and observed a 40-fold range of fold-changes in expression (**Figure 2D**). We determined the fold-change of variants by normalizing induced and uninduced measurements to negative controls and then calculated the ratio of normalized induced expression to normalized uninduced expression. Promoters consisting of the consensus −10 and −35 sites exhibited the highest fold-changes, however, these values were highly variable depending on the variant’s operator site composition (**Figure 2E**). Amongst promoters containing these core sites, we found that operators in the *proximal* site were largely deterministic of fold-change, with promoters containing strong operators (O_1_ and O_sym_) in the *proximal* site yielding 4.61-fold higher fold-changes on average than promoters containing weak operators in the *proximal* site (*p* = 1.44 × 10^−6^, Welch’s t-test). We attribute this to the importance of the downstream operator in blocking RNAP binding and transcriptional initiation^10,46^. As expected, promoters containing O_sym_ in the *proximal* site generally drove the highest fold-change, however, pairing with another O_sym_ in the *distal* site surprisingly resulted in decreased fold-change relative to other variants. Notably, while the consensus core promoter containing O_sym_ in both the *proximal* and *distal* sites yielded a fold-change of 4.63x, its counterpart containing the weaker O_1_ variant in the *proximal* site drove a superior fold-change of 8.97x. Upon deeper investigation, we found that although this promoter containing O_sym_ in both the *proximal* and *distal* sites had 1.77-fold lower uninduced expression compared to its counterpart containing the weaker O_1_ in the *proximal* site, its induced expression was also 3.43-fold lower **(Figure S3A)**. Thus, having O_sym_ in both the *proximal* and *distal* sites decreased expression in the induced state by a larger magnitude than in the uninduced state, resulting in lower fold-change. This demonstrates that simply using the strongest binding elements available may not yield the highest fold-change levels.

### Biophysical modeling of inducible promoter activity

We set out to elucidate this important point by combining our experimental measurements with a statistical mechanical model of binding to clarify under what conditions optimal fold-change can be achieved. To validate our conceptual understanding of how the promoter and operator elements collectively give rise to gene expression within our promoter architecture, we developed a statistical mechanical model of binding that could analyze the thousands of promoter combinations to extrapolate the behavior of a general promoter and clarify under what conditions optimal fold-change can be achieved. Moreover, this model could further validate our experimental finding that combining the strongest RNAP sites (the consensus −35 and −10) with the strongest LacI sites (a *proximal* and *distal* O_sym_) did not yield the largest fold-change. One possible explanation for this observation is that even at 1 mM IPTG, a small number of active LacI will still be active^47^. While the amount of time any repressor is expected to be active is small, the large binding affinity to O_sym_ sites may nevertheless enable measurable repression^48,49^.

To that end, we modeled this promoter architecture by enumerating the various promoter states containing all combinations of RNAP binding, LacI binding, and LacI looping (**Figure S4A**). We assume that all states where RNAP is bound and the proximal LacI site is not bound give rise to gene expression r_max_, whereas all other states have a small background level of gene expression r_min_^9,50^. The relative probability of each state is given by *e*^−β*E*^ where *E* equals the sum of all binding free energies arising from binding or looping (**Figure S4A**). Upon summing the contributions from all states, the average gene expression is given by:

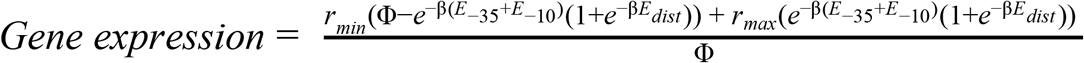

where

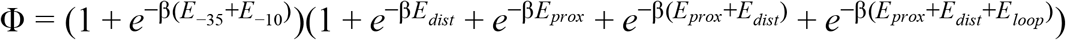

represents the partition function (the sum of the Boltzmann weights for all states). This compact form signifies that gene expression only arises when RNAP is bound (and contributes *E*_−35_ + *E*_−10_ to the free energy) and the distal LacI site is either unoccupied or occupied (adding free energy 0 or *E*_*dist*_, respectively). We used this form of gene expression to infer the binding energies of each promoter element and compared the resulting fits for the 1,493 different promoters in absence of IPTG (**Figure 3A**, parameter values in **Figure S4B**). Moreover, this model enables us to extrapolate the gene expression for promoter architectures with arbitrary binding strengths spanning the theoretical parameter space **(Figure 3B)**.

With only slight modification, the above equation for gene expression can also be used to model these same promoters at 1 mM IPTG. In the absence of IPTG, all repressors are in the active state, in which they are capable of binding the promoter^47^. When 1 mM IPTG is added, only a small fraction, 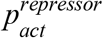, of these repressors will be active. Hence the Boltzmann weights *e*^−β*E*_*dist*_^ and *e*^−β*E*_*prox*_^ of bound LacI, which are proportional to the number of active repressors, must all be multiplied by 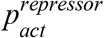. This can be achieved by modifying:

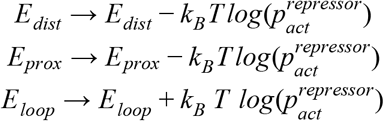

in the equation for gene expression above, where the last relation for the looping free energy arises because looping corrects for the effective concentration of a singly bound repressor binding with its other dimer. In summary, by introducing the single additional parameter 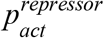, we can extend our characterization of the 1,493 promoters in the absence of IP aTlsGotoinclude their gene expression at 1 mM IPTG.

**Figure 3.**
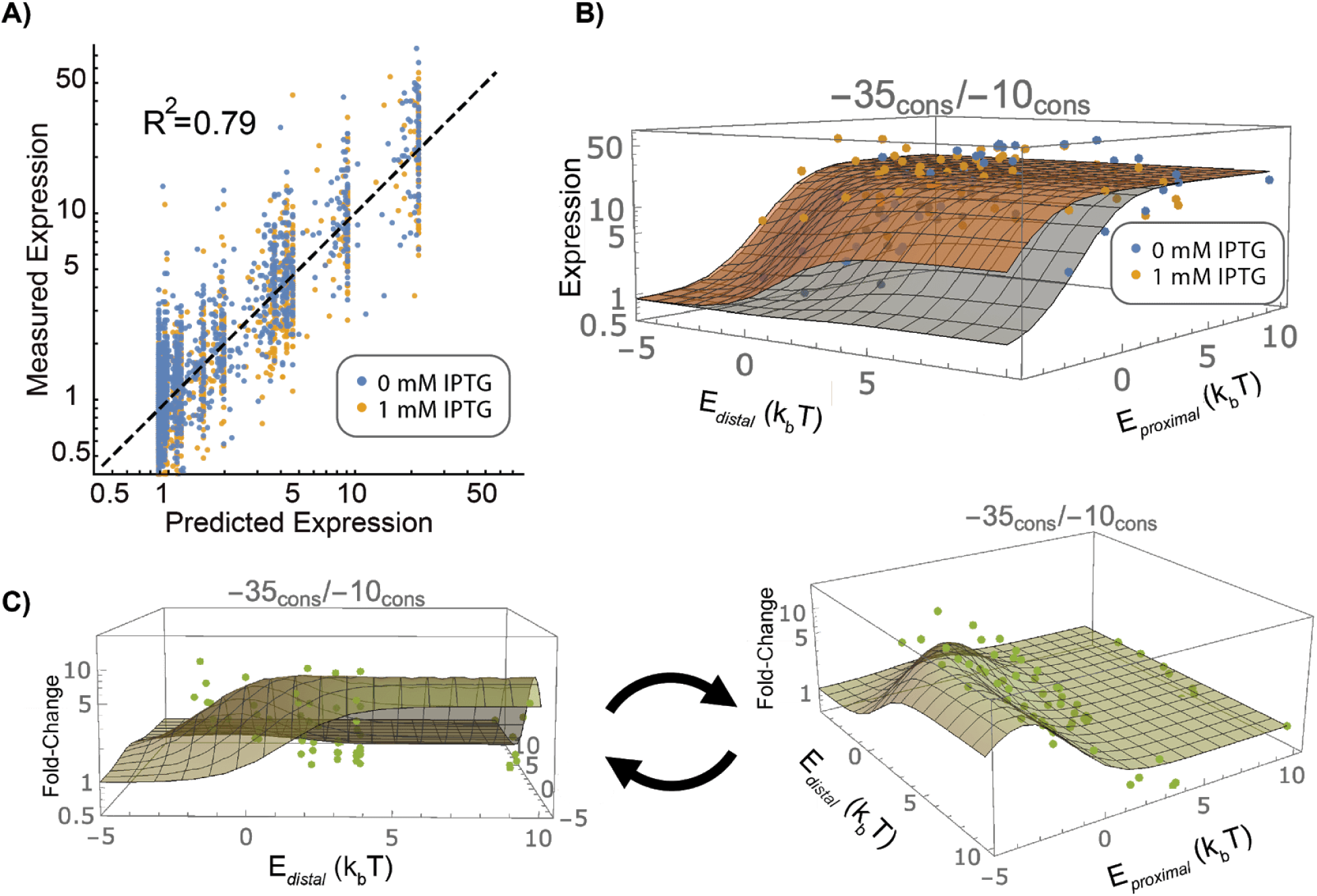
Thermodynamic modeling of lacUV5 promoter architecture. **A)** Correlation between actual *lacUV5* variant expression and expression fit by our thermodynamic model (*R*^2^ = 0.79, p < 2.2 × 10^−16^). **B)** Induced and uninduced gene expression across the *distal* and *proximal* site binding energy parameter space. **C)** Fold-change (FC) in gene expression as a function of *distal* and *proximal* binding site energies. In Panels B and C, each dot represents experimental data whereas the grid lines denotes the inferred expression of a promoter with the proximal (E_*proxima*l_) and distal (E_*distal*_) LacI binding energy shown.

**Figure 3A,B** shows that the fit gene expression aligns with our experimental measurements using the value 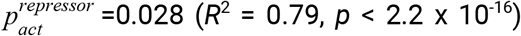. This implies that 28 of every 1000 repressor molecules are in the active state at 1 mM IPTG, or equivalently that each repressor fluctuates sporadically between an active and inactive state but will on average only spend 2.8% of the time in the active state. We note that this value for the fraction of active repressors inferred from our data is 28 times larger than a previously imputed value for 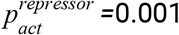 at 1 mM IPTG^47^. Enforcing this previous value for 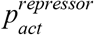 while fitting the model resulted in comparable parameter values (**Figure S4C**) and overall fit (*R*^2^ = .79, *p* < 2.2 × 10^−16^).

Given the gene expression in the presence and absence of IPTG, we could now explore how the fold-change depends upon the binding energy of the two LacI sites. Returning to our earlier result, we confirmed that using the consensus −35/−10 RNAP binding site together with a proximal and distal O_sym_ (binding energy − 2.4*k_B_T*; **Figure S4B**) for LacI leads to suboptimal fold-change **(Figure 3C)**. As we hypothesized, these repressor binding sites are sufficiently strong to overcome the small amount of active repressors per cell, leading to reduced gene expression even at 1 mM IPTG **(Figure 3B)**. Instead, the promoter architecture that maximizes fold-change couples the strong −10 and −35 RNAP elements with near-maximal LacI operator site strengths that are sufficiently strong enough to repress in the absence of IPTG but not in the presence of saturating IPTG.

### Alternative IPTG-inducible promoter architectures exhibit different behaviors in induced and uninduced conditions

We next explored whether alternative promoter architectures could exhibit higher fold-change and reduced leakiness compared to the canonical *lacUV5* architecture. Previous work has demonstrated that varying the architecture, or the positioning of operator sites, within inducible systems can alter the input-output response to IPTG^14,51^. However, we lack the understanding of promoter sequence-function relationships necessary to systematically design novel promoter architecture variants with desirable behaviors. Instead, we leveraged our multiplexed screening approach to engineer novel inducible promoter architectures. By screening large numbers of variants, we explored the sequence space around three rationally-designed architectures, allowing us to identify desirable variants as well as learn how binding site strengths relate to activity in each design.

#### Additional operator sites can promote or antagonize induction response

Based on our previous characterization of the 1,600 Pcombo variants, we speculated whether an additional distal operator site could improve the fold-change of promoters. In particular, we expected that an additional distal site would enhance repression by increasing the likelihood of loop formation. To investigate this, we synthesized and tested 2,000 *lacUV5* variants within a library we call Pmultiple. This library resembled Pcombo except for the inclusion of an additional modular LacI binding site, which we refer to as the ‘*distal+*’ site, immediately upstream of the *distal* binding site. The final design was composed of each combination of five *distal+* operator sites, five *distal* operator sites, four −10 elements, four − 35 elements, and five *proximal* operator sites for a total of 2,000 variants **(Figure 4A, top)**. Using our MPRA, we measured expression for 1,638 of these variants (81.9%) in the absence of IPTG and at 1 mM IPTG with an average of 8-9 barcodes measured per variant **(Figure S2B)**. To determine the effect of the *distal+* site in this architecture, we compared the fold-change of each Pmultiple variant to Pcombo variants composed of the same *distal*, −35, −10, and *proximal* sites. We limited our analysis to studying promoters with consensus core promoter elements as well as an O_1_ or O_sym_ *proximal* site to best capture the repressive effects of the *distal+* element. We found that the effect of adding the *distal+* site to the Pcombo architecture spanned a 5.4-fold range where it would either increase or decrease fold-change of the variant depending on the identity of the *distal* site **(Figure 4A, bottom)**. We observed that a strong *distal+* operator site is consistently able to compensate for a weak *distal* operator site to decrease leakiness and improve fold-change. The greatest effect was observed when adding an O_1_ *distal+* site to the variants with the weakest *distal* operator, O_3_, resulting in a 2.93-fold increase in fold-change. However, when the *distal* site was already strong, adding a *distal+* operator actually decreased the fold-change in expression. Upon further investigation, we found that in these cases where a strong *distal* site was already present, the addition of a strong *distal+* site actually increased leakiness and induced expression of the system, suggesting that the *distal+* site may be inhibiting repression of the promoter by the *distal* site **(Figure S5A,S5B)**. Thus, we conclude that additional *distal* operator sites can improve the fold-change of an inducible systems by reducing the uninduced expression, however, they can have negative effects if coupled with an already strong *distal* site.

**Figure 4.**
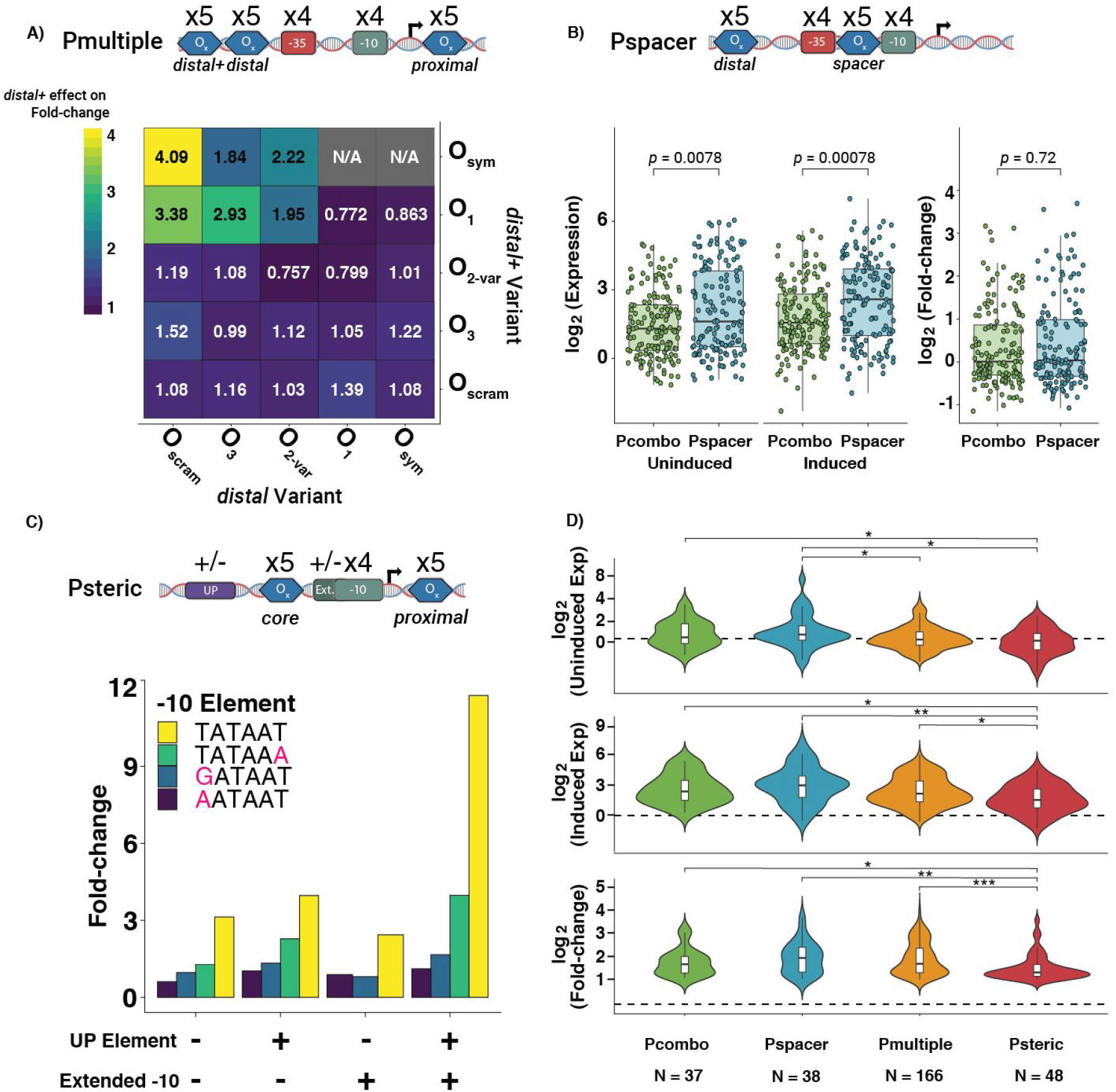
Optimizing alternative IPTG-inducible promoter architectures. **A) Top:** Design for Pmultiple library. **Bottom:** The average effect of the *distal+* site (rows) on fold-change given the *distal* site (column). Here we examine consensus −10/−35 promoters containing O_1_ or O_sym_ in the *proximal* site. **B) Top:** Design for Pspacer library. **Bottom:** Comparison of uninduced expression, induced expression, and fold-change between variants composed of the same sequence elements in the Pspacer and Pcombo architectures (Two-sided Mann-Whitney U-tests). We examined only active promoters containing a consensus −10 and/or −35 sequence. **C) Top:** Psteric library design. **Bottom:** The fold-change of promoters containing O_1_ in both the *core* and *proximal* sites and a 56 bp inter-operator distance. Here we examine the effect of the −10 element in conjunction with the strongest UP and extended −10 element combinations. **D)** Distributions of uninduced expression, induced expression, and fold-change for variants with log_2_(fold-change) > 0 in each library. Dashed line separates active from inactive sequences and is set as the median of the negative controls + 2*median absolute deviation (Two-sided Mann-Whitney U-tests with Benjamini-Hochberg correction, ‘*’ = *p* < 0.05, ‘**’ = *p* < 0.01, ‘***’ = *p* < 0.001).

#### Repositioning operator sites alters activity independent of sequence element composition

Next, we explored how the repositioning of operator sites influences repression of the *lacUV5* promoter. A previous work indicated that operator sites placed within the spacer region, the segment of DNA between the −10 and −35 elements, enabled strong repression^13^. Notably, this positions the operator such that it directly competes with RNAP binding. Furthermore, this architecture is desirable for synthetic applications as it avoids placing operators within coding sequences and 5’ UTRs, which is the case for promoters with operators at the *proximal* site^14^. To explore this concept in depth, we synthesized Pspacer, a library of 4,400 variants containing all combinations of five *distal* operator sites, four −35 elements, four −10 elements, and five *spacer* operator sites **(Figure 4B, top)**. Because this *spacer* region is 17 bp and the LacI operators we use are 21 bp, operator sequences were truncated by 2 bp at their termini so as not to overlap the −10 and −35 motifs. In order to determine an optimal spacing between the *distal* and *spacer* operator sites, we also tested these combinations with inter-operator distances between 46 and 56 bp. We recovered expression data for 3,769 (85.7%) of these variants in the absence of IPTG and at 1 mM IPTG with an average of 7 barcodes per variant **(Figure S2C)**. The distance between the *spacer* and *distal* operator sites did not appear to significantly affect the fold-change of the promoters at the *p* < 0.05 threshold (ANOVA), which may be because some of the tested distances were insufficient to enable the formation of DNA loops^17,43^ **(Figure S6A, S6B)**.

With all operator spacings tested appearing equivalent, we subset our analysis to variants with an inter-operator distance of 55 bp, which is reportedly amenable to looping^43^. Similarly to variants with the Pcombo architecture, we only observed strong induced expression with promoters containing −10 and −35 elements resembling the consensus (**Figure S6C**). To see how this change in architecture altered the performance of these promoters, we compared Pspacer variants to respective Pcombo promoters composed of the same sequence elements. We observed that promoters with the Pspacer architecture had on average 2.16-fold higher uninduced and 1.93-fold higher induced expression **(Figure 4B, bottom)**. This may be because fewer repressed states are possible in this architecture, thereby pushing the system to be more active. Alternatively, this increased expression may be due to greater spacer %AT content within *spacer* LacI sites^20,52^(**Table S3**). Despite these higher expression values, Pspacer variants had comparable levels of fold-change to corresponding variants of the Pcombo architecture (**Figure 4B, bottom**).

#### Altering RNAP binding contacts

Finally, we explored whether altering the RNAP contacts could modify the behavior of inducible systems. Although all promoters we have tested thus far were designed to contact the RNAP through the σ70 −35 and −10 hexamer elements, previous reports have suggested that it may be possible to engineer promoters lacking −35 elements^53,54^. In these cases, it is proposed that additional compensatory binding sites for transcription factors or the RNAP are necessary to recruit RNAP and enable transcription. In addition to the −35 and −10 motifs, RNAP binding may be enhanced by an extended −10 TGn^55,56^ motif as well as an AT-rich UP element^57,58^ upstream of the −35 that may stabilize RNAP through contacts with the α-subunit. However, it has not been directly shown that these additional points of contact are sufficient to compensate for the lack of a −35 element.

We synthesized and tested a library of 1,600 *lacUV5* variants called Psteric containing each combination of four −10 elements, five *core* operator sites centered at −26 in place of the −35 element, five *proximal* operator sites, and four UP elements in the presence or absence of an extended −10 motif (**Figure 4C, top**). Furthermore, we positioned the *proximal* operator site centered at either the canonical +11 position or at the +30 position. At the +30 position, the *proximal* operator is 56 nucleotides away from the *core* operator, which is near an optimal distance for repression loop formation^29^. We recovered expression data for 1,369 of these variants (85.6%) in the absence of IPTG and at 1 mM IPTG with an average of 8 barcodes per variant (**Figure S2D**). We first examined library variants lacking functional LacI operator sites to see whether any combinations of −10 elements, extended −10 elements, and UP elements could yield functional promoters. Although weak or no transcription was detected from promoters with only a −10, we found that relatively strong promoters could be created by the addition of an UP-element and extended −10, with up to 13-fold greater expression than promoters containing a consensus −10 but lacking both these supplementary elements **(Figure S7A)**. Interestingly, these sites had little effect on their own and we could only detect a significant effect when both UP element and extended −10 motifs were added. Thus, we found that promoters designed with this architecture could be engineered.

Next, we explored which designs in this library could enable inducible promoter systems. First, we find that the variants with the highest fold-change were constructed with *proximal* operator sites located at the +30 position relative to the TSS (**Figure S7B**). Secondly, we find that the success of this architecture relies on the presence of an UP element, an extended −10, and a strong −10 motif. When all three of these elements are present, promoters of this architecture appear to exhibit up to a 11.8-fold response to IPTG **(Figure 4C, bottom)**. Despite the apparent viability of this architecture, we found that the highest expressing promoters generally contained O_scram_ or O_1_ *core* operator sites (**Figure S7C**). In these cases, we found that the operator sites contained partial matches to the −35 motif, although these were not in the optimal position relative to the −10 motif (**Figure S7D**). However, several comparatively weak but functional variants were recovered that do not appear to contain any cryptic −35 elements (**Figure S7C**), thereby demonstrating that it is possible to engineer inducible promoters lacking this key motif.

### Comparison of optimized alternative *lacUV5* promoter architectures

To gauge how each of our alternative promoter architectures perform relative to one another, we compared the distributions of fold-changes between variants in each library. To focus our analysis on inducible variants, we limited our analysis to the variants in each library with fold-change > 2. Of the thousands of promoters tested with each architecture, relatively few were capable of induction, highlighting the surprising difficulty in engineering these systems. We find that each architecture generated promoters with similarly wide ranges of uninduced expression, induced expression, and fold-changes (**Figure 4D, Table S8**). However, overall comparisons revealed significant differences in the properties of these distinct architectures. In particular, Pcombo variants exhibited significantly higher uninduced expression compared to Psteric variants, while Pspacer variants exhibited significantly higher uninduced expression compared to Pmultiple and Psteric variants (*p* < 0.05, Two-sided Mann-Whitney U-test with Benjamini-Hochberg correction). Additionally, Psteric variants exhibited significantly lower fold-changes and induced expression compared to Pcombo, Pmultiple, and Pspacer variants (*p* < 0.04, Two-sided Mann-Whitney U-test with Benjamini-Hochberg correction).

### Validation of functional inducible variants using a fluorescent reporter

Ultimately, we sought to identify inducible promoter variants within each library that were superior to the canonical *lacUV5* promoter. From each of the four libraries, we identified promoter sequences exhibiting high fold-change with low leakiness and individually integrated these variants into the *essQ-cspB* locus of *E. coli* MG1655. To evaluate the fold-change in expression of these promoters, we used flow cytometry to measure expression of a sfGFP reporter gene at both the uninduced (0 mM IPTG) and fully induced state (1 mM IPTG) (**Figure 5**). All variants exhibited improved fold-change compared to *lacUV5* (min: 9.5x, max: 21.0x, *lacUV5*: 4.1x) and a majority displayed distinct separation between the uninduced and induced states. In particular, variants of Pmultiple exhibited fold-changes more than 5-fold higher than *lacUV5*. Many variants, especially the Psteric promoters, exhibited low leakiness while maintaining comparable induced expression. Activity measurements using flow cytometry were well-correlated with MPRA measurements (Induced: *r* = 0.701, Uninduced: *r* = 0.981, Fold-change: *r* = 0.885) (**Figure S8)**. Lastly, we wanted to determine whether these architectures resulted in different input-output relationships in response to IPTG induction. To test this, we measured GFP expression of these eight *lacUV5* variants at six IPTG concentrations in triplicate spanning 3 orders of magnitude (0, 0.001, 0.005, 0.01, 0.1, 1 mM) using a Tecan plate reader (**Figure S9**). We observe that the fold-change across all eight promoters followed similar trends, beginning induction at similar concentrations of IPTG (0.01 mM IPTG) and generally saturating at 1 mM IPTG. In all cases, variants exhibited far greater levels of fold-change compared to *lacUV5.* Overall, our multiplexed exploration of rationally design variants allowed us to engineer novel promoters with reduced leakiness and higher fold-changes compared to *lacUV5*.

**Figure 5.**
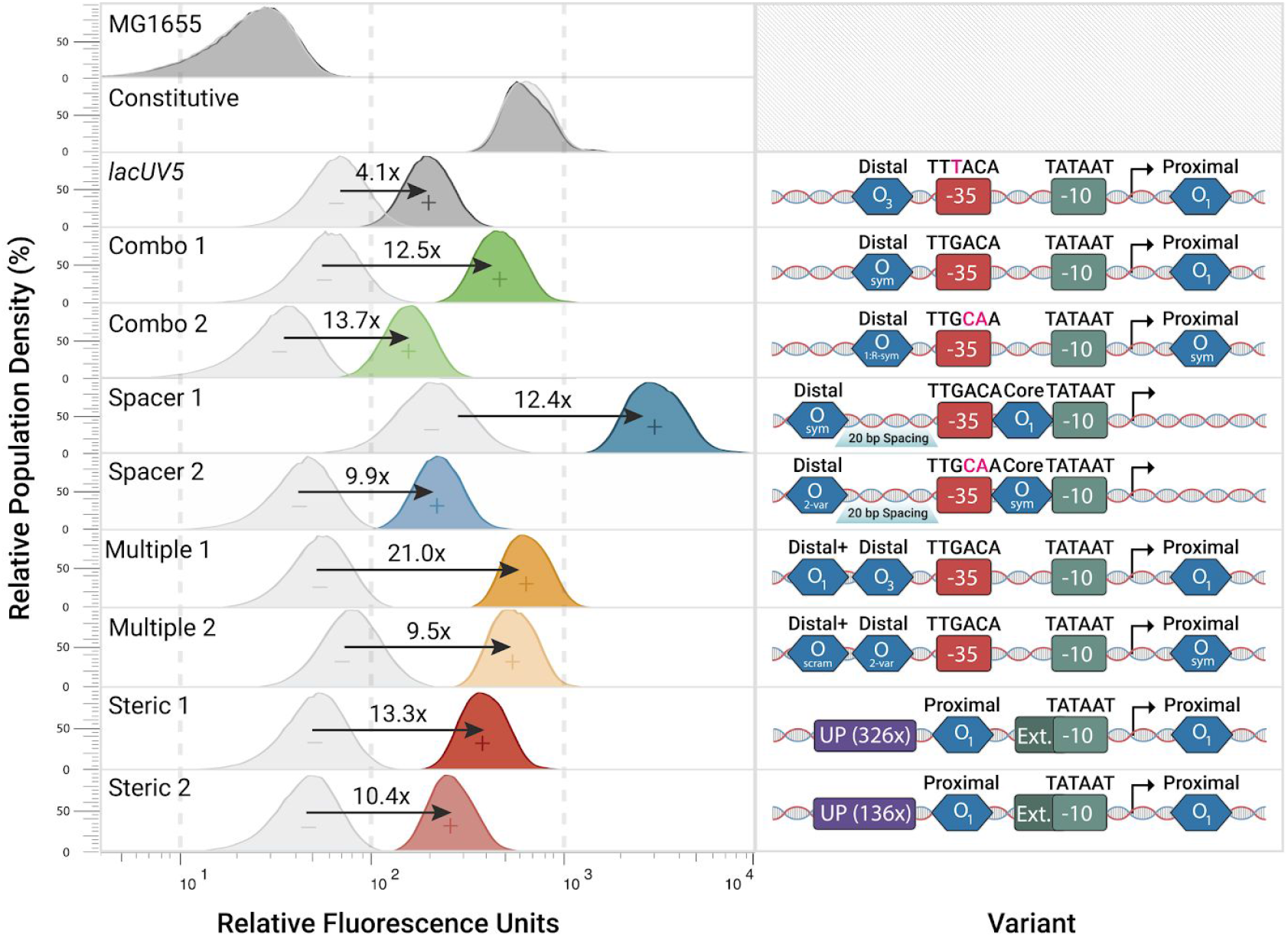
Characterization of functional inducible variants using a fluorescent reporter. Fluorescence measurements of selected variants for induced and uninduced states determined using flow cytometry. Fold-change of each variant was estimated after background subtracting induced and uninduced expression. “−” represents the promoter in uninduced state while “+” represents induction after 1 mM IPTG.

## Discussion

While current methods for tuning inducible systems involve arbitrarily manipulating operator sites and core promoter elements, these approaches tend to provide little insight into the combinatorial interactions modulating expression. Here, we implemented a MPRA to measure gene expression of nearly 9,000 different promoter variants to learn the design rules for multiple P*lacZYA* sequence architectures. We find that the ability of the canonical *lac* promoter to induce largely depends on a complex interplay between the repressor sites and the core promoter. Notably, RNA polymerase and a repressor compete for binding, such that promoters containing near-consensus −35 and −10 σ70 elements are functionally irrepressible unless matched with correspondingly strong repressor sites. However, as in previous studies, we observe that the strongest repressor sites may even repress in the presence of inducer, reducing the fold-change in gene expression. We developed a thermodynamic model for this architecture which demonstrated how sufficiently strong LacI binding sites can reduce gene expression even in the presence of saturating inducer. Both the model and our empirical measurements agree that large fold-change is achieved by using strong RNAP binding elements (e.g. the consensus −35 and −10 sequences) together with the strongest O_sym_ operator in the distal site and the second strongest O_1_ operator in the proximal site.

Beyond studying interactions between elements within the canonical *lacZYA* promoter architecture, our investigation of different promoter architectures revealed striking interactions between repressor sites and core promoter elements, uncovering several characteristics of optimal promoters. In our study of the Pmultiple architecture we found that adding an upstream *distal+* repressor site could compensate for weaker *distal* sites, however, if an already strong *distal* site was present, addition of a *distal+* site would inhibit repression of the promoter through an unknown mechanism. Secondly, we showed that repositioning repressor sites in the Pspacer architecture increased overall uninduced and induced expression levels relative to equivalent promoters in the Pcombo architecture, however, this did not alter the overall fold-change of the promoters. Lastly, our studies of the Psteric architecture show that it is possible to engineer promoters lacking −35 elements, although these promoters require UP, extended −10, and near-consensus −10 elements to be functional.

Ultimately, this systems analysis of inducible promoter regulation demonstrates the utility of combining rational design with large-scale multiplexed assays. Testing sequence libraries in multiplexed formats enabled exploration of a wide distribution of functional designs as well as the discovery of promoter variants with desirable properties. Additionally, this high-throughput assay provides a reliable means for exploring the effects of variation across distant regions of the functional sequence landscape, which can reveal novel insights into promoter mechanisms and sequence-function relationships.

## Acknowledgements

This work was supported by the National Science Foundation Graduate Research Fellowship 2015210106 to G.U., National Institutes of Health New Innovator Award DP2GM114829 to S.K., Searle Scholars Program to S.K., U.S. Department of Energy (DE-FC02-02ER63421 to S.K.), UCLA, and Linda and Fred Wudl. We thank the UCLA BSCRC high throughput sequencing core and Technology Center for Genomics and Bioinformatics for technical assistance; All past and present members of the Kosuri lab for technical feedback; Suzannah Beeler for thoughtful discussions; and Reid C. Johnson for manuscript feedback. Lastly, we thank the UCLA Molecular Biology Interdepartmental Graduate Program and UCLA Bioinformatics Interdepartmental Graduate Program.

## Author contributions

T.C.Y., G.U., W.L.L., J.E.D., J.S., G.B., T.E., and S.K. designed the study. T.C.Y. and K.D.I generated the sequence libraries. T.C.Y., M.B., W.L.L., J.S., and G.B. performed the experiments. T.C.Y., G.U., J.E.D., and T.E. analyzed the data. W.L.L. designed the figures. T.E. developed the statistical mechanics model. T.C.Y., G.U., W.L.L., J.E.D., J.S., G.B., and T.E. wrote the manuscript. All authors editing and approved the manuscript.

## Competing interests

The authors declare no competing interests.

## MATERIALS AND METHODS

### Promoter Library Design

#### Transcription factor spacing

A library of 624 variants were created to test the effects of altering spacing between LacI, AraC, GalR, GlpR, LldR, and PurR operator sites. The core promoter PlacL8-UV5, is the endogenous lacZYA promoter region with L8 and L29 mutations in the CAP site to render it catabolite insensitive (−55 C->T, −66G->A) as well as UV5 mutations in the −10 region to increase activity (−9,−8 GT->AA)^59–61^. Pairs of 23 bp operator sites were acquired from endogenous loci reported by RegulonDB^41^ (ver 8.0) (**Table S1**). For sites under 23 bp in length, the surrounding sequence of the native genomic context was included. In all cases, the downstream site found at the endogenous loci, with respect to the regulated promoter orientation, was used as a *proximal* site in our designs while the upstream sequence was used as the *distal* site. For each pair of operator sites, a series of variants were designed where the *proximal* operator was centered at +12 (spanning +1 to +23) and the *distal* operator varied from positions −83 to −116. Similar series of variants were also designed in which the sequence of the *proximal* site or *distal* site was shuffled to obviate activity of the operator.

#### Pcombo

A library of 1,600 *lacUV5* variants composed of each combination of 10 *proximal* operator sites, 10 *distal* operator sites, four −10 elements, and four −35 elements was designed. The operator sites were selected to span a wide range of lacI binding affinities (**Table S2**). These consisted of two native LacI operators (O_1_ and O_3_) and a variant of the native O_2_ lac operator with three mutations (O_2-var_). Additionally, O_sym_ and six other synthetic operators (O_1:R-sym_, O_2:L-sym_, O_2:R-sym_, O_3:L-sym_, O_3:R-sym_) were used with the latter being designed by creating palindromic sequences based on either the left or right halves of each native sequence. Lastly, a scrambled operator (O_scram_) composed of a random scrambling of the O_1_ sequence served as a negative control. The −10 and −35 sites were selected to span a range of binding affinities for RNA Polymerase and obtained from a previous characterization^6,8,20^ (**Table S4, S5**). Each variant was composed of a combination of these elements placed onto catabolite insensitive (L8, L29 mutant), *lacZYA* promoter with the *proximal* site placed at +11 and the *distal* site placed at −90, which was found to enable strong loopinging in the assay of transcription factor spacing.

#### Pmultiple

A library of 2,000 *lacUV5* variants composed of each combination of one of five *distal+* operator sites, five *distal* operator sites, five *proximal* operator sites, four −10 elements, and four −35 elements was designed. The O_1_, O_3_, O_2-var_, O_sym_, and O_scram_ operators from the Pcombo library were selected as the five operator sites for testing. Additionally, the same −10 and −35 elements from the Pcombo library were selected. This library was constructed with sequence elements placed in the same positions as the Pcombo library, with the exception of the *distal+* sequence being placed immediately upstream of the *distal* site.

#### Pspacer

A library of 4,400 *lacUV5* variants composed of each combination of five *distal* operator sites, four −35 elements, four −10 elements, and five *spacer* operator sites was designed. In order to fit the 17-bp spacer region, two base pairs were trimmed from each end of the *spacer* operator sites (**Table S2**). The same operators, −10 elements, and −35 elements from the Pmultiple library were selected. Lastly, the distal operator site was tested at 10 different spacings relative to the core promoter, ranging from 20-30 bp from the 5’ most end of the −35 element. These 20-30 bp spacings resulted in inter-operator distance of 46-56 bp.

#### Psteric

A library of 800 *lacUV5* variants composed of each combination of four −10 elements, five *core* LacI sites centered at −26, five *proximal* operator sites, and one of four UP elements in the presence or absence of an extended −10 motif was designed. The same operator sites and −10 elements from the Pmultiple library were selected. *Proximal* operator sites were and tested when centered at both the +11 and +30 positions relative to the TSS. The UP elements selected were obtained from a previous characterization and range in their abilities to enhance transcription^20,62^ (**Table S6**). Additionally, the extended −10 element TGG was used as this is the most commonly found version of an extended −10^56^.

### Library Cloning

The library was synthesized by Agilent and then resuspended in 100 uL of elution buffer before cloning into plasmid pLibacceptorV2 (Addgene ID no. 106250). The transcription factor spacing library was ordered separate from the other libraries, which were altogether synthesized and tested in a multiplexed pool. First, the library was amplified with KAPA SYBR FAST qPCR Master Mix (#KK4600) utilizing primers GU 132 and GU 133 at 10 uM to determine Cq values. Afterwards, the library was amplified with NEBNext^®^ Q5^®^ Hot Start HiFi PCR Master Mix (#M0543S) at 11 cycles using primers GU 132 and GU 133 as well, in triplicate. Replicates were pooled, then cleaned with Zymo Clean and Concentrator Kit (#D40140).

To barcode the library, each library was amplified with NEBNext^®^ Q5^®^ Hot Start HiFi PCR Master Mix (#M0543S) for 10 cycles using primers GU 132 and GU 134. Library ends were then digested with SbfI-HF (NEB #R3642S) and XhoI (NEB #R0146S) by incubating at 37°C for 1.5 hours. The plasmid vector, pLibAcceptorV2, was first maxi-prepped with QIAGEN Plasmid Maxi Kit (#12162), concentrated with a Promega Wizard SV Gel and PCR Clean-up System (#A9281), and digested with SbfI-HF (NEB #R3642S), SalI-HF (NEB #R3138S), and rSAP (NEB #M0371S) for 1.5 hours at 37°C. Insert (library) and vector (pLibAcceptorV2) were ligated using T7 DNA Ligase (NEB #M0318S), incubating at room temperature for 1 hour. The plasmid was then transformed into DH5α electrocompetent *E. coli* cells (New England Biolabs C2989K) and plated for 24 hours at 30°C on LB + kanamycin (25 ug/mL) agar plates. These plates were then harvested in 5 mL of LB and 400×10^6^ cells (based on OD_600_) were grown overnight in 450 mL LB + kanamycin (25 ug/mL). This plasmid, consisting of the library cloned into pLibacceptorV2, was isolated and concentrated in the same method as described earlier.

To clone RiboJ::sfGFP into the plasmid, RiboJ::sfGFP was first amplified with NEBNext^®^ Q5^®^ Hot Start HiFi PCR Master Mix (#M0543S) for 25 cycles using primers GU 99 and GU 100 at 10 uM. This amplicon was then digested with BsaI-HF (NEB # R3535) and NcoI-HF (NEB #R3193S) for 1.5 hours at 37°C. pLib was digested with BsaI-HF (NEB # R3535) and NheI (NEB# R3131S). pLib vector was then ligated with the GFP insert using T7 DNA Ligase (NEB #M0318S), incubating at room temperature for 1 hour. This plasmid was next transformed into DH5α electrocompetent cells and plated for 24 hours of growth at 30°C as well, yielding “pLib_sfGFP” plasmid after maxi-prep.

### Library Integration

The pLib_sfGFP plasmid was first digested with SalI-HF (NEB#R3138S) and NheI (NEB# R3131S) to remove background. This was then transformed into the landing pad strain, an engineered^20^ *E. coli* MG1655 derivative (Yale Coli Genetic Stock Center no. 6300), and grown overnight for 24 hours at 30°C. The following day, plates were scraped and 800 million cells in 200 mL of LB + kan (25 ug/mL) was inoculated overnight at 30°C.

For library integration, glycerol stocks of landing pad strain with the integration plasmid were grown overnight in 200 mL + kan (25 ug/mL) at 30°C. 200 million cells from this overnight culture was inoculated the next day into 250 mL LB + 0.2% arabinose + 25 ug/ml Kan at 30°C for 24 hours to induce recombination. The following day, 800 million cells of induced overnight was inoculated into 80 mL LB + 25 ug/mL Kan at 42°C for heat cure. This was grown to log phase (OD 0.3-0.7) for about 1.5 hours. 200 million cells from this log phase culture were plated at 42°C for 16 hours in undiluted, 10^−5^, and 10^−6^ dilutions. Plates grown overnight were then scraped, and 400 million cells inoculated into 200 mL LB + Kan 25 ug/mL for overnight growth at 37°C. Ultimately, this was plated again at 30°C to validate integration (GFP instead of mCherry) and then glycerol stocked after colony PCR for further confirmation.

### Barcode Mapping

The promoter and barcode region from pLib was prepared for sequencing and downstream mapping of the barcodes to their respective variants. Two PCRs were performed to prepare pLib samples for sequencing, the first of which adds sites for the sequencing primer whereas the second PCR adds the adaptors for Illumina sequencing and a unique index DNA label. Each barcode mapping was performed in duplicate.

For the first PCR, the library was amplified with KAPA SYBR FAST qPCR Master Mix (#KK4600) with primers GU 60 and GU 79 at 5 uM to determine Cq values. Afterwards, the library was amplified with NEBNext^®^ Q5^®^ Hot Start HiFi PCR Master Mix (#M0543S) at 11 cycles using primers GU 60 and GU 79 at 5 uM as well in triplicate. Replicates were pooled, then cleaned with Zymo Clean and Concentrator Kit (#D40140), eluting into 10 uL of Ultra-pure H2O.

For the second PCR, illumina adapters P7, P5, and a unique DNA index were added. The product from the first PCR was amplified with primers GU 70 and GU 86 at 5 uM to determine Cq values. Afterwards, the library was amplified with NEBNext^®^ Q5^®^ Hot Start HiFi PCR Master Mix (#M0543S) at 10 cycles using primers GU 70 and GU 86 at 5 uM. Since different primers add different indices to each sample, we re-ran the second PCR with a different set of primers to serve as redundancy and allow us to compare sequencing replicates. This process was repeated in a separate PCR, with primers GU 70 and GU 87 also at 5 uM.

Ultimately, each technical replicate was performed in duplicate, cleaned with Zymo Clean and Concentrator Kit (#D40140), and ran on a 1.0% agarose gel for final confirmation. After quality assessment, samples were sequenced on an Illumina Nextseq 500 using a Paired end 300-cycle kit (2×150 bp). Barcodes were mapped to their respective promoter variants using the pipeline from Urtecho et al. 2018^20^. In brief, paired-end reads are merged using PEAR^63^ (version 0.9.1). We then extract the first 150 bp of each read, which encodes the promoter variant, as well as the last 20 bp encoding the barcode and generate a list of barcode-variant associations. Finally, we perform additional filtering steps for quality control purposes.

### Library Growth and Sequencing Preparation

Library pellets were prepared in both Induced and Uninduced conditions. First, glycerol stocks were inoculated in 100 mL of MOPS with 0.2% glucose + kanamycin (25 ug/mL) at 30°C for 16 hours overnight. The following day, the overnight culture was diluted to OD 0.0005, inoculated into 200 mL MOPS + kanamycin (25 ug/mL) with 0.2% glucose, and grown at 37°C to OD 0.5-0.55 (about 5 hours) both with 1 mM IPTG and without.

To harvest RNA pellets, the culture was first cooled for two minutes in an ice slurry while periodically swirling. For each sample, three 50 mL aliquots of culture were poured into pre-chilled tubes and spun for two minutes at 13000xg at 4°C. The supernatant was poured off. RNA was extracted from *E. coli* pellets using Qiagen RNEasy Midiprep kit (#75142). We performed technical replicates of this extraction (separate RNA extractions of the same culture) with the operator spacing library and biological replicates (Different cultures grown in parallel before separately extracting). Subsequent wash steps concentrated isolated RNA with Qiagen Minelute Cleanup Kit (#74204). Next, isolated RNA was converted to cDNA with Thermo Fisher SuperScript IV (#18090010) following manufacturers directions.

To harvest gDNA pellets, 5 mL samples of each culture were then spun down for four min @ 5000xg. Supernatant was then poured out. DNA from each pellet was then isolated with Zymo Research ZR Plasmid Miniprep Kit (#D4015) for use as normalization.

The barcoded cDNA was amplified with NEBNext^®^ Q5^®^ Hot Start HiFi PCR Master Mix (#M0543S) from 1 ug of gDNA for 14 cycles with primers GU 59 and GU 60 at 5uM. The product was cleaned with Zymo Clean and Concentrator Kit (#D40140). 1 ng of this sample was amplified again for 10 cycles with primers GU 65-68 and GU 70 for indexing, yielding 8 total samples; technical replicates for induced and uninduced cDNA, and induced and uninduced gDNA. Both prepared DNA and RNA library samples were quantified with Agilent Tapestation, then sent for sequencing on HiSeq2500 (SE 50-cycle) to the *Broad Stem Cell Research Center at UCLA*.

### Data processing

Following RNA-Seq and DNA-Seq of the barcodes, we quantify the relative abundance of each barcode. Demultiplexed RNA and DNA reads for each biological replicate were converted to counts of each barcode via a custom bash script that extracts barcode sequences from individual reads and counts the number of observed reads for each barcode. These barcode counts were normalized using the following formula:

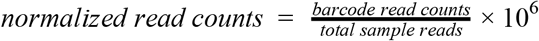

Normalized read counts were then merged by common barcode to yield a comprehensive data frame containing normalized read counts for each barcode in each replicate. This dataframe was then merged with the barcode mapping data to map normalized read counts to their corresponding promoter. Multiple barcodes could map to a single promoter, thereby providing replicability, and any promoter that contained fewer than 3 barcodes in any sample were removed. After this filtering step, promoter expression for each replicate was calculated using the following formula:

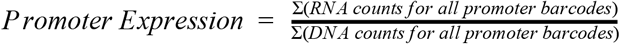

To normalize promoter expression between induced and uninduced samples, expression of each promoter was normalized to the median negative control promoter expression in its respective biological replicate. Lastly, the mean expression of the biological replicates was calculated to obtain final expression values for the induced and uninduced conditions.

### Thermodynamic model of gene expression

For the Pcombo library, initial guesses for the binding energies of each LacI operator site were used as inputs and refined when fitting a statistical mechanics model to the Pcombo promoter expression data. The coefficient of determination (*R*^2^) between fit and actual gene expression values was calculated using log_10_-transformed values to reduce the effects of large expression outliers.

### Code Availability

The Mathematica notebook used for the thermodynamic model as well as all code for recreating plots are available at: https://github.com/timcyu/inducible_architecture.

### Individual Promoter Variant Cloning

Two promoters were selected from each of the libraries, yielding eight total promoters in addition to two controls (a constitutive promoter and UV5). Individual promoter variants were selected from our library of variants based on the highest fold-change (Induced over uninduced expression) and fold-change:noise ratio (fold-change over uninduced expression). These sequences were ordered from IDT as gBlocks^®^ Gene Fragments. Full RiboJ:sfGFP was PCR isolated from the original library. Since promoters were to be measured individually, we did not include a barcode in synthesis. Plasmid vector, pLibacceptorV2 was linearized with SbfI-HF (NEB #R3642S) and SalI-HF (NEB #R3138S).

After synthesis by IDT, promoters were amplified using primers GU 142, GU 89, and NEBNext^®^ Q5^®^ Hot Start HiFi PCR Master Mix (#M0543S). Each reporter was assembled with Gibson Assembly^®^ Master Mix (NEB #E2611S) using 30 bp overlaps between the plasmid pLibAcceptorV2, the promoter, and RiboJ:sfGFP. Each assembled reporter was separately transformed into *E. coli* DH5α Chemically Competent *E. coli* (NEB #C2987H) yielding 10 total transformed *E. coli* strains containing their respective promoter, RiboJ:sfGFP, and Kanamycin antibiotic resistance. Afterwards, the promoter and downstream GFP segment were sequenced from isolated colonies using the same set of primers, GU 142 and GU89, to confirm correct constructs. All products were cleaned with Zymo Clean and Concentrator Kit (#D40140) except for pLibAcceptorV2, which was cleaned with Promega Wizard SV Gel and PCR Clean-up System (#A9281) after DNA isolation with QIAGEN Plasmid Maxi Kit (#12162).

### Individual Promoter Variant Integration

*E. coli* strains containing library members were grown overnight for 16 hours in 5 mL of Luria Broth and kanamycin (25 mg/uL). Afterwards, the plasmid was isolated using Zymo ZR Plasmid Miniprep Kit (#D4054) formed into an electrocompetent MG1655 containing an engineered landing pad within the *essQ-cspB* intergenic locus^20^ and plated on LB and kanamycin (25 μg/mL) at 30°C. Two colonies per promoter were resuspended in LB, and inoculated into 5 mL of LB + kanamycin (25 μg/mL) for overnight growth.

Each promoter was separately integrated into the *essQ-cspB* locus using Cre-Lox mediated cassette exchange. Following overnight growth, cells of this culture were inoculated into 5 mL of LB, kanamycin (25 μg/mL), and 0.2% arabinose (g/mL) and grown for 24 hours to induce integration of the reporter cassette. After integration of the reporter cassette through the arabinose-induced Cre system, residual plasmid was removed through heat-curing. 200 million cells were inoculated into 3 mL of LB and kanamycin (25 μg/mL) and grown at 42°C for about 1.5 hours to reach log phase (OD 0.3-0.7). After this growth, cells were diluted to 10^-4 and plated on LB + kanamycin (25 ug/mL) plates overnight at 42°C to complete the heat-curing process.

### Plate Reader Assay

Glycerol stocks for each promoter were scraped and inoculated into liquid cultures containing MOPS EZ-Rich Media (TEKNOVA #M2105) and 25 ug/mL of kanamycin at 30°C for overnight growth in 5 mL disposable culture tubes. The following day, each promoter was diluted to OD 0.005 in 500 uL of MOPS EZ-Rich Media (TEKNOVA #M2105) with 0.2% glucose (g/mL) and 25 ug/mL of kanamycin and set up for plate reader analysis in triplicates across an IPTG gradient: 0, 0.001, 0.005, 0.01, 0.1, 1 mM. After samples were grown for five hours at 37°C, 100uL aliquots were transferred into 96-Well Flat Bottom Microplates. Measurements were taken for wavelengths 650 nm (measures OD) and 520 nm (measures GFP) on the Tecan Infinite M1000 Pro No.30064852 plate reader. Data was analyzed in Excel with the four reads per time point per well averaged and divided by the OD measurement to calculate the GFP fluorescence.

### Flow Cytometry

Glycerol stocks for each promoter were first scraped and inoculated into liquid cultures containing MOPS EZ-Rich Media (TEKNOVA #M2105) and 25 ug/mL of kanamycin at 30°C for overnight growth. The following day, cells grown overnight were diluted to an OD of 0.002 in MOPS EZ-Rich Media (TEKNOVA #M2105) with 0.2% glucose (g/mL) and 25 ug/mL of kanamycin at 30°C. These cells were then transferred to 100 mL flasks all containing 15 mL of MOPS EZ-Rich Media + 0.2% glucose. 1 mM IPTG + 25 ug/mL kanamycin were added to the “Induced” cultures whereas 25 ug/mL kanamycin was added to the “Uninduced” cultures. These cultures were then grown at 37°C for 3.5 hours. 5 mL of each sample was spun down, the supernatant was decanted, and the cell pellets were resuspended in 1 mL PBS (GIBCO^®^ PBS Phosphate-Buffered Saline 10010023). 1 mL of each sample was filtered into a Falcon 5 mL Polystyrene Round-Bottom Tube with Cell-Strainer Cap. *E. coli* MG1655 was used as a negative control for GFP expression while a constitutively active library member was used as positive. Data was collected using a BioRad S3 Cell Sorter and analyzed in FlowJo (version 10.0.8r1). Fold-change was calculated by dividing the median GFP fluorescence of the induced samples by the median fluorescence of the induced samples.

