## Supplementary Figures S1-S9, Tables S1-S8 for "Multiplexed characterization of rationally designed promoter architectures deconstructs combinatorial logic for IPTG-inducible systems"

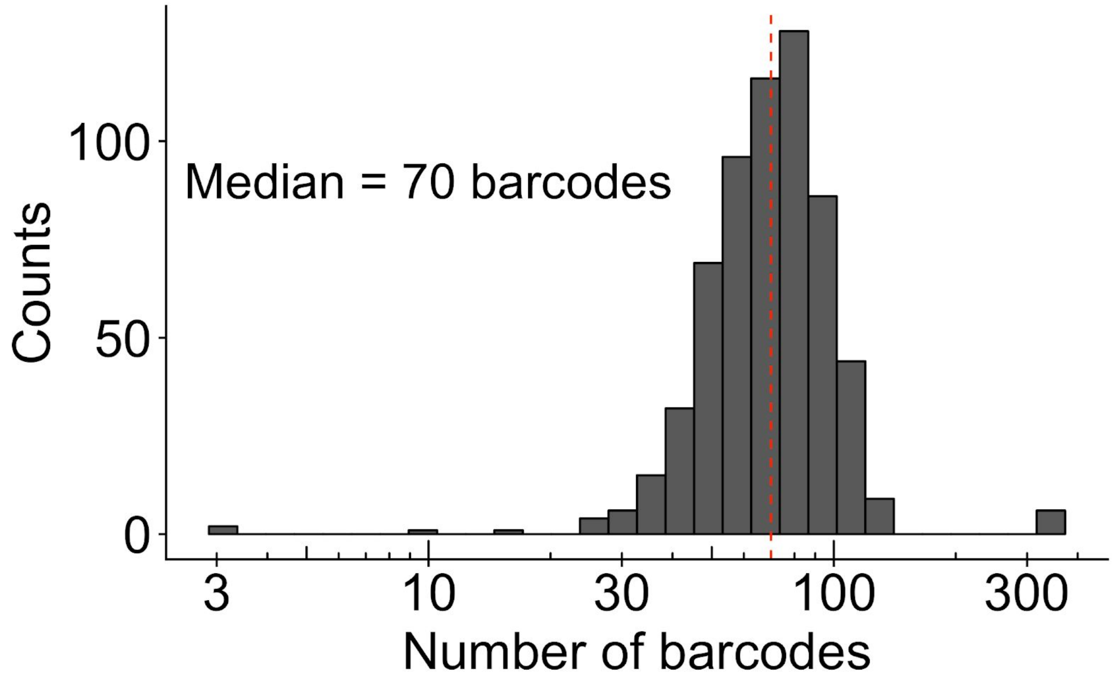


**Figure S1) Distribution of the number of unique barcodes for operator spacing variants.** We recovered promoter activity data for 615 of the 624 (98.6%) variants in the operator spacing library. On average, we measured the expression of 70 unique barcodes per variant.

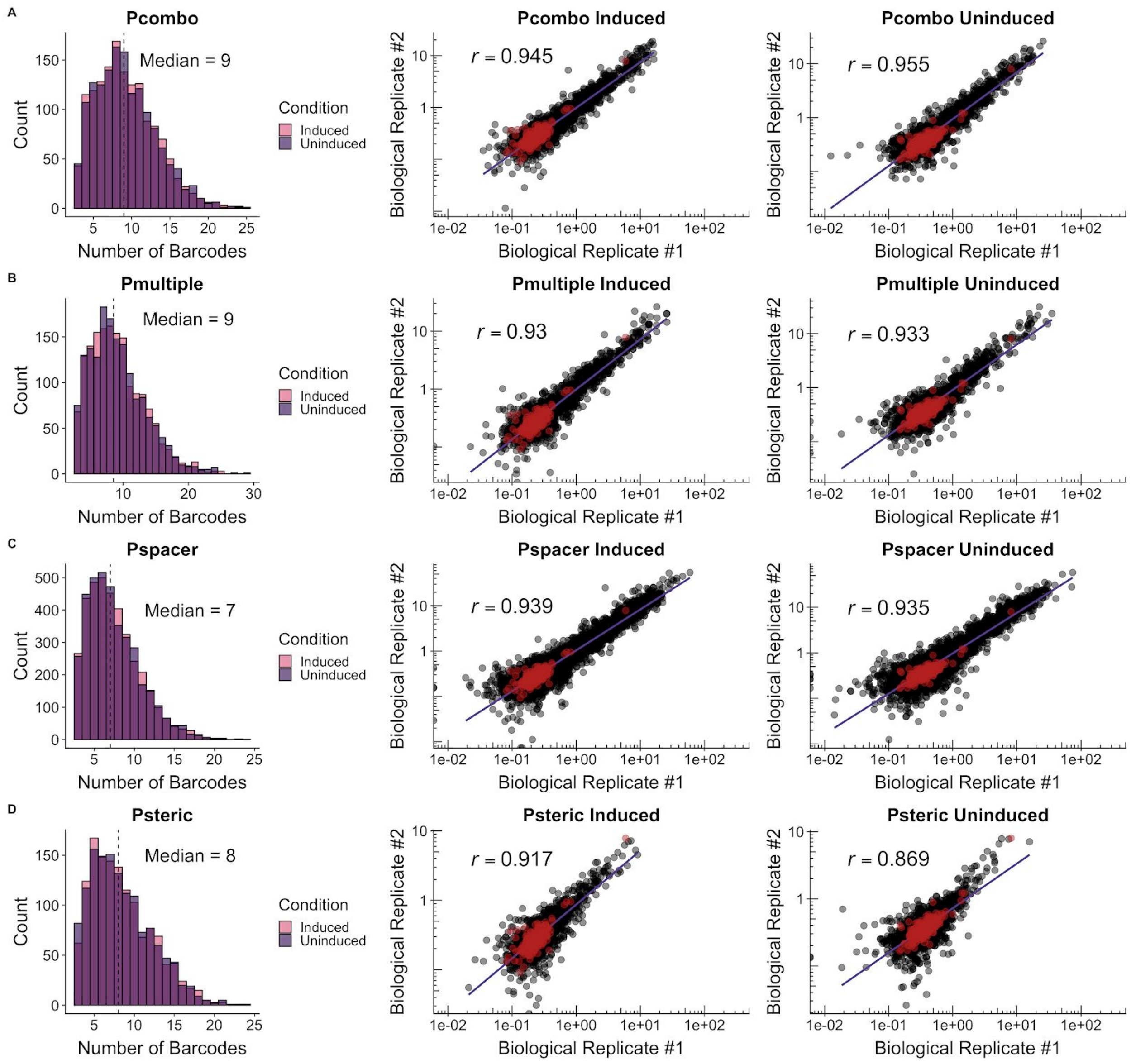


**Figure S2) Distributions of unique barcode sequences and correlations between biological replicates.** Library quality statistics for **A)** Pcombo **B)** Pmultiple **C)** Pspacer **D)** Psteric. All variants in each library show strong correlation between biological replicates ($p < 2.2 \times 10^{-16}$).

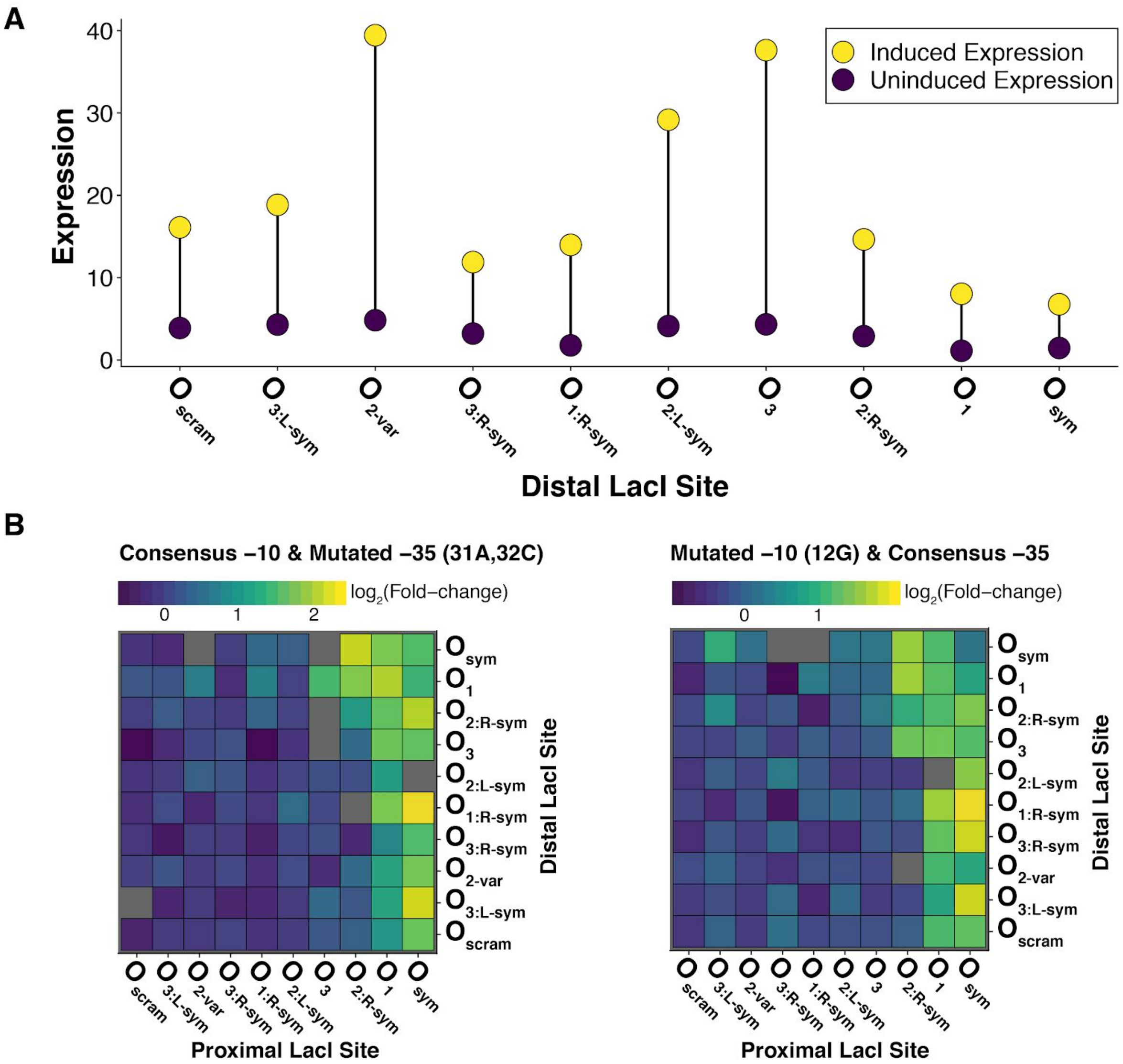


**Figure S3) Optimal repressor binding sites in the Pcombo library is conditional based on the identity of the core promoter. A)** Expression ranges of variants containing consensus -10, consensus -35, and *proximal* $O_{sym}$. Promoters containing $O_{sym}$ in both the *proximal* and *distal* sites have weaker induced expression compared with promoters containing *proximal* $O_{sym}$ and a weaker *distal* site. **B)** Fold-change for Pcombo variants containing one of the consensus -10/-35 elements coupled with a near consensus -10/-35 element. The best operator combination by fold-change differs when either the -10 or -35 element is mutated, suggesting an interplay between the repressor sites and core promoter strength.

A)

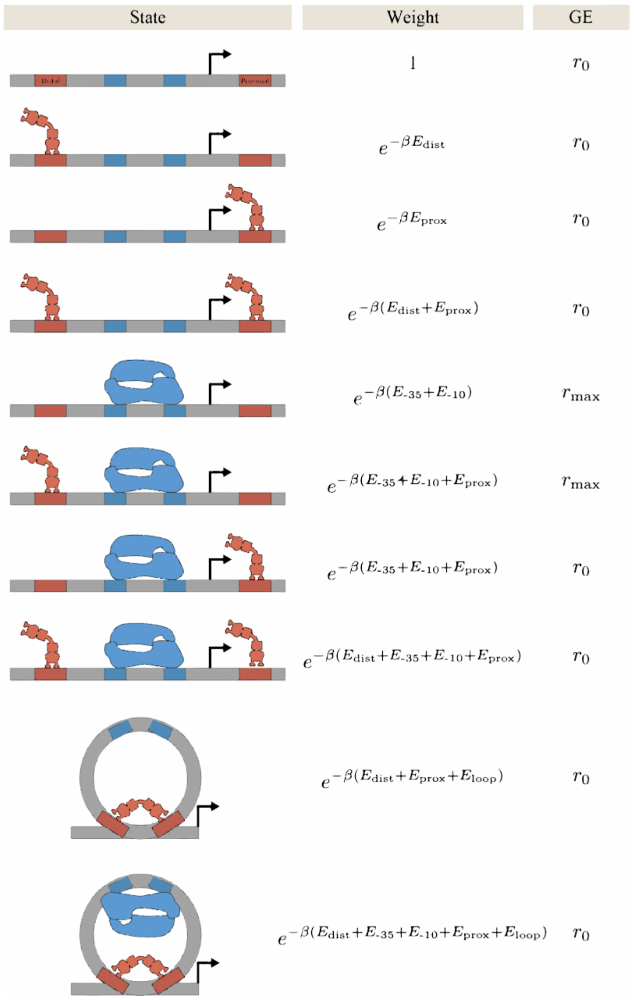


B)

| Parameter | Value |
|---|---|
| $r_{min}$ | 0.879 |
| $r_{max}$ | 25.14 |
| $E_{loop}$ | −2.49 |
| $p_{act}^{repressor}$ | 0.0282 |
| $E[lac_{O1}]$ | −0.23 |
| $E[lac_{O1}^{rightSym}]$ | 3.01 |
| $E[lac_{O2}]$ | 3.91 |
| $E[lac_{O2}^{leftSym}]$ | 27.88 |
| $E[lac_{O2}^{rightSym}]$ | 1.58 |
| $E[lac_{O3}]$ | 2.01 |
| $E[lac_{O3}^{leftSym}]$ | 3.76 |
| $E[lac_{O3}^{rightSym}]$ | 25.96 |
| $E[lac_{Oscram}]$ | ∞ |
| $E[lac_{Osym}]$ | −2.09 |
| $E[-10_{cons}]$ | 0. |
| $E[-10_{7A}]$ | 4. |
| $E[-10_{12A}]$ | 5.03 |
| $E[-10_{12G}]$ | 3.77 |
| $E[-35_{cons}]$ | −1.96 |
| $E[-35_{30C\_33T}]$ | 3.58 |
| $E[-35_{31A\_32C}]$ | 0.66 |
| $E[-35_{33T}]$ | 1.7 |

C)

| Parameter | Value |
|---|---|
| $r_{min}$ | 0.89 |
| $r_{max}$ | 24.09 |
| $E_{loop}$ | −4.66 |
| $p_{act}^{repressor}$ | 0.001 |
| $E[lac_{O1}]$ | −0.23 |
| $E[lac_{O1}^{rightSym}]$ | 4.68 |
| $E[lac_{O2}]$ | 5.87 |
| $E[lac_{O2}^{leftSym}]$ | ∞ |
| $E[lac_{O2}^{rightSym}]$ | 3.23 |
| $E[lac_{O3}]$ | 3.65 |
| $E[lac_{O3}^{leftSym}]$ | 5.61 |
| $E[lac_{O3}^{rightSym}]$ | ∞ |
| $E[lac_{Oscram}]$ | ∞ |
| $E[lac_{Osym}]$ | −2.21 |
| $E[-10_{cons}]$ | 0. |
| $E[-10_{7A}]$ | 4.1 |
| $E[-10_{12A}]$ | 5.14 |
| $E[-10_{12G}]$ | 3.87 |
| $E[-35_{cons}]$ | −2.06 |
| $E[-35_{30C\_33T}]$ | 3.6 |
| $E[-35_{31A\_32C}]$ | 0.64 |
| $E[-35_{33T}]$ | 1.7 |

**Figure S4) A thermodynamic model for *lacUV5*. A)** Thermodynamic states of *lacUV5* architecture and their corresponding Boltzmann weights. The probability that the system is in each state is proportional to the relative values of the Boltzmann weights. The system is assumed to elicit a background level of gene expression (GE) given by $r_{min}$ unless RNAP is bound with no repressor bound to the *proximal* site, in which case a larger level of promoter activity $r_{max}$ is evoked. **B)** Best-fit parameter values inferred by fitting this model to the 1,600 promoters with this architecture, simultaneously considering their gene expression with and without 1mM IPTG. **C)** Best-fit parameter values inferred when $p_{act}^{repressor}$ at 1 mM IPTG is manually set to its previously reported value 0.001.

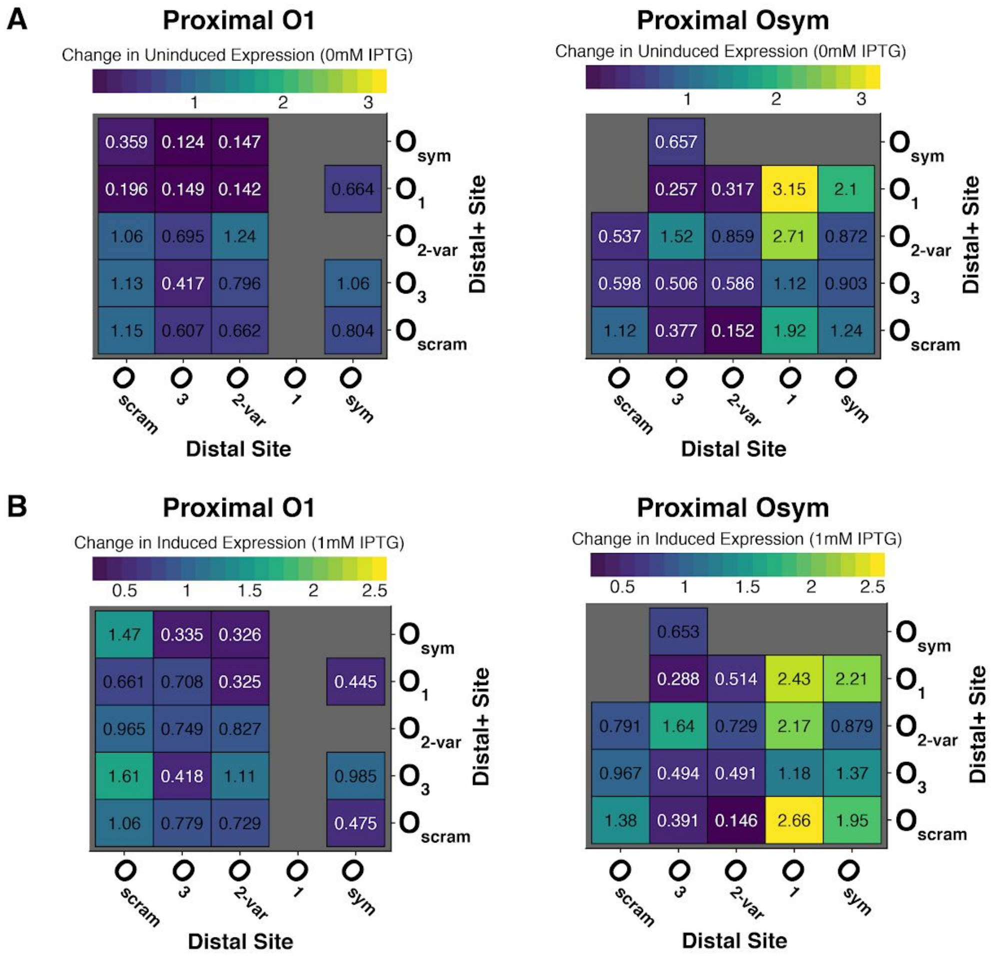


**Figure S5) Effect of *distal*+ site on induced and uninduced expression in the Pmultiple library. A)** Change in uninduced expression of Pmultiple variants with *proximal* $O_1$ (left) and $O_{sym}$ (right) relative to their Pcombo counterparts. **B)** Change in induced expression of Pmultiple variants with *proximal* $O_1$ (left) and $O_{sym}$ (right) relative to their Pcombo counterparts.

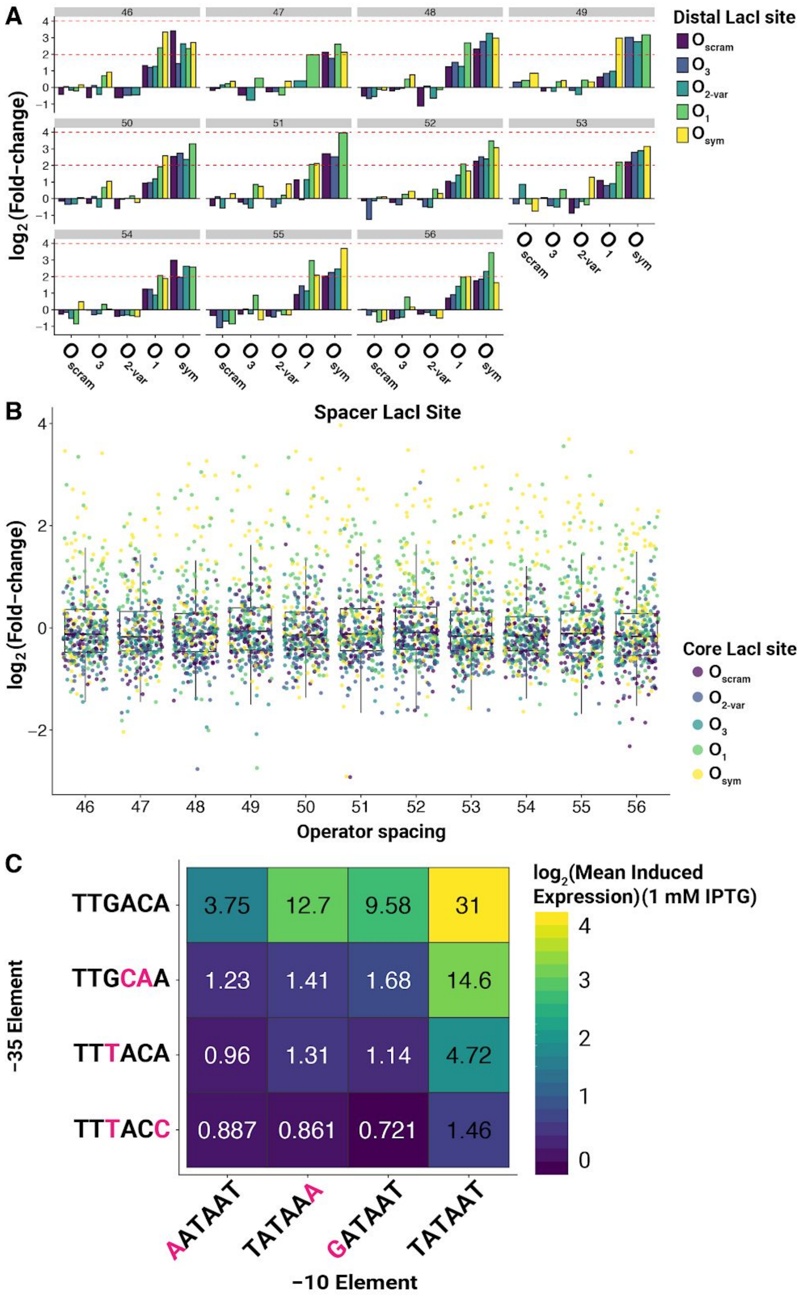


**Figure S6) Fold-induction and induced expression are modulated by the strengths of repressor sites and the identity of the core promoter in the Pspacer library. A)** Fold-change of *spacer* and *distal* combinations at each operator spacing for Pspacer variants containing the consensus core promoter. An inter-operator distance of 55 bp yielded consistent and strong repression across all combinations of functional operators. **B)** Distribution of fold-changes for each distance between operator sites show little effect due to operator distance. **C)** Mean induced expression for each combination of -10 and -35 combinations amongst Pspacer library variants.

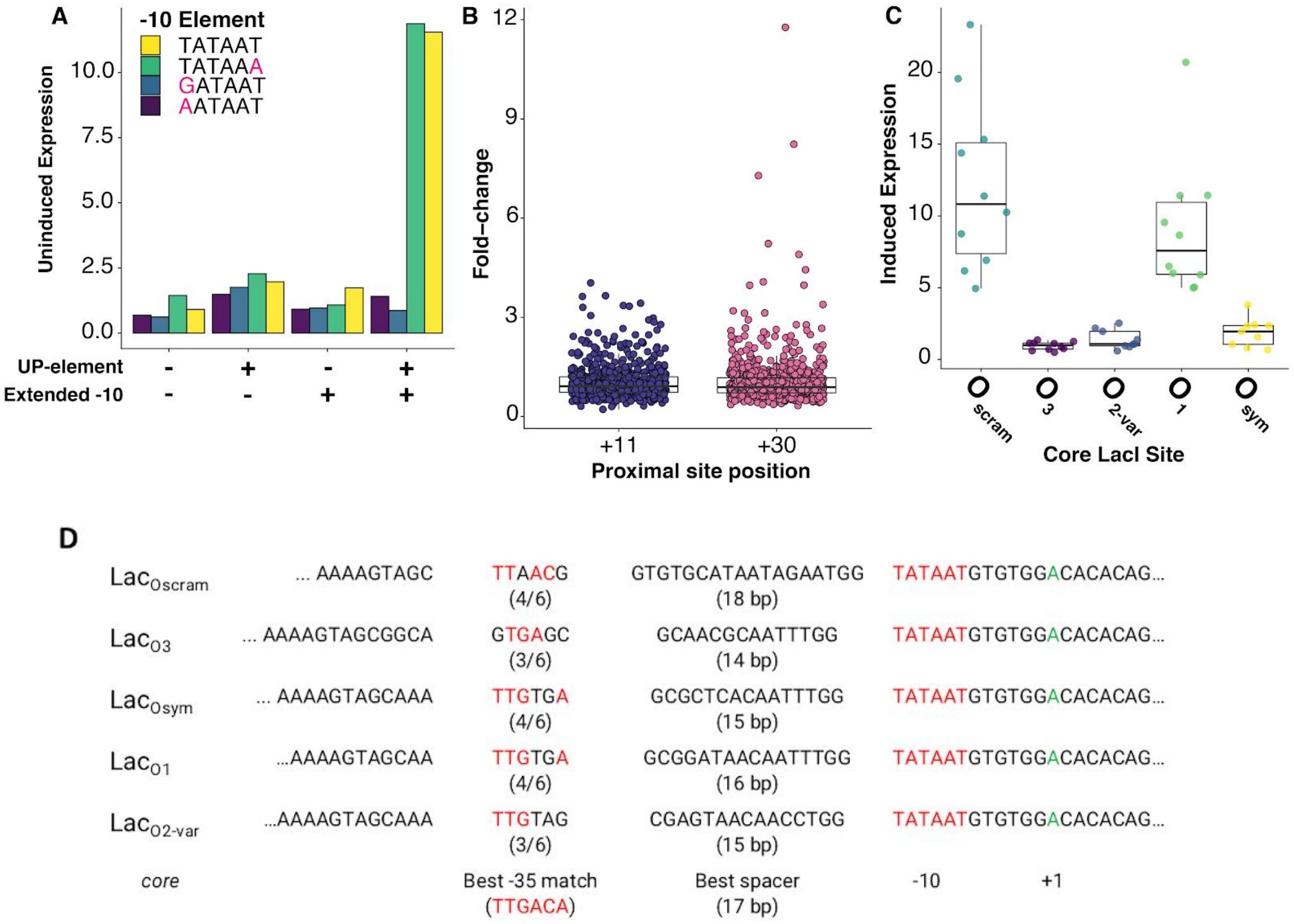


**Figure S7) Operator site distance and composition influence Psteric architecture viability. A)** Uninduced expression for promoters containing *proximal* and *core* Oscram sites, split by combination of UP element and extended -10 motif. **B)** Psteric variants with high fold-change are only observed when containing a *proximal* site centered at +30, resulting in a 56 bp spacing between operator sites. **C)** Strongly expressed Psteric variants primarily contained *core* operator sites containing partial matches to the -35 motif, despite not being in the optimal position relative to the -10 motif. **D)** Potential -35 motif sequence and spacing for each *core* operator site. LacOscram and LacO1 operator sites contain 4/6 bases matching the -35 motif, although these matches are positioned at non-optimal spacer lengths.

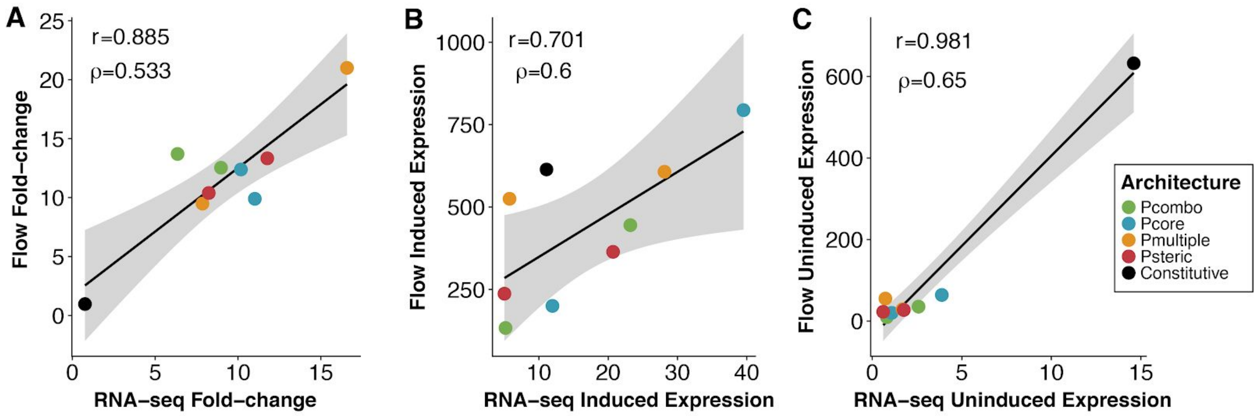


**Figure S8) Correlations between flow cytometry and RNA-seq. A-C)** Comparison of RNA-Seq and flow cytometry measurements for promoters individually characterized in **Figure 5.** Strong Pearson correlations are reported between fold-change ($r$ = 0.885, $p$ = 0.001), induced expression ($r$ = 0.701, $p$ = 0.03), and uninduced expression ($r$ = 0.981, $p$ = 3.3 x $10^{-6}$) measurements for flow cytometry and RNA-seq. Moderately strong Spearman correlations are also reported between fold-change (ρ = 0.533, $p$ = 0.15), induced expression (ρ = 0.6, $p$ = 0.10), and uninduced (ρ = 0.65, $p$ = 0.07) measurements for flow cytometry and RNA-seq.

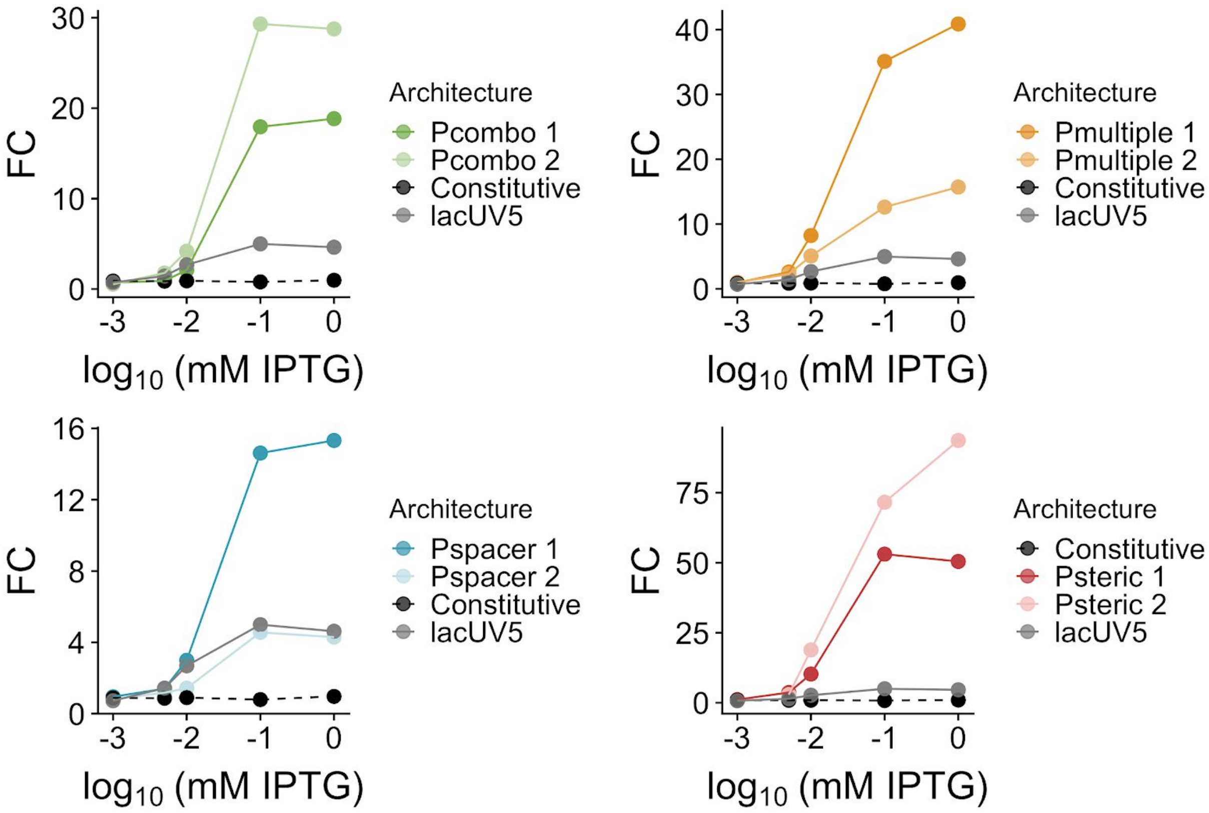


**Figure S9) Architecture input-output relationships following IPTG induction.** Input-output response to IPTG for variants from each architecture compared to *lacUV5* and a constitutively active variant (Constitutive).

| TF | *Distal* site (5'>3') | *Proximal* site (5'>3') | *Source Operon* |
|---|---|---|---|
| LacI | GGGCAGTGAGCGCAACGCAATTA | GAATTGTGAGCGGATAACAATTT | *lacZYA* |
| GalR | TCTTGTGTAAACGATTCCACTAA | TACCGGTGGTAGCGGTTACATTG | *galETKM* |
| AraC | GAAGAAACCAATTGTCCATATTG | CCATAGCATTTTTATCCATAAGA | *araBAD* |
| PurR | GTTGAGGAAAACGATTGGCTGAA | TTTAAGCAAACGGTGATTTTGAA | *purA* |
| GlpR | AAAATGTTCAAAATGACGCATGA | AAATGGTAAAAAACGAACTTCA G | *glpTQ* |
| LldR | AAGAATTGGCCCTACCAATTCTT | CACAATTGGCAGTGCCACTTTTA | *lldPRD* |

**Supplementary Table 1) Transcription factor binding sites to test operator spacings.** Sequences were acquired from RegulonDB[40].

| Operator Variant | Sequence(5'>3') | Inferred Binding energy ($K_bT$) |
|---|---|---|
| $O_1$ | AATTGTGAGCGGATAACAATT | -0.4 |
| $O_{2\text{-var}}$ | AAATTGTAGCGAGTAACAACC | 2.9 |
| $O_3$ | GGCAGTGAGCGCAACGCAATT | 2.0 |
| $O_{sym}$ | AAATTGTGAGCGCTCACAATT | -2.4 |
| $O_{scram}$ | TTAACGGTGTGCATAATAGAA | ∞ |
| $O1_{rightsym}$ | AATTGTTATCGGATAACAATT | 2.8 |
| $O2_{rightsym}$ | AAATGTGAGCCGCTCACATTT | 1.4 |
| $O2_{leftsym}$ | GGTTGTTACTCAGTAACAACC | 4.7 |
| $O3_{rightsym}$ | AATTGCGTTGGCAACGCAATT | 3.9 |
| $O3_{leftsym}$ | GGCAGTGAGCGGCTCACTGCC | 3.2 |
| $O_1$-spacer | TTGTGAGCGGATAACAA | --- |
| $O_{2\text{-var}}$-spacer | ATTGTAGCGAGTAACAA | --- |
| $O_3$-spacer | CAGTGAGCGCAACGCAA | --- |
| $O_{sym}$-spacer | ATTGTGAGCGCTCACAA | --- |

| | | |
|---|---|---|
| $O_{scram}$-spacer | AACGGTGTGCATAATAG | --- |

**Supplementary Table 2) Operator site sequences and binding energies.** Operator sequences used to assemble libraries.

| Operator Variant | Sequence (5'>3') | %AT-content |
|---|---|---|
| O1-spacer | TTGTGAGCGGATAACAA | 58.8% |
| O2-var-spacer | ATTGTAGCGAGTAACAA | 64.7% |
| O3-spacer | CAGTGAGCGCAACGCAA | 58.8% |
| Osym-spacer | ATTGTGAGCGCTCACAA | 52.9% |
| Oscram-spacer | AACGGTGTGCATAATAG | 58.8% |
| WT lacUV5 spacer | TTTATGCTTCCGGCTCG | 47.1% |

**Supplementary Table 3) Pspacer operator sites and %AT content.**

| -35 element name | Sequence (5'>3') |
|---|---|
| minus35cons | TTGACA |
| minus35_31A_32C | TTGCAA |
| minus35_33T | TTTACA |
| minus35_30C_33T | TTTACC |

**Supplementary Table 4) -35 element sequences used in this work.** These -35 sequences for RNAP recognition were previously reported and result in a wide range of binding affinities.

| -10 element name | Sequence (5'>3') |
|---|---|
| minus10cons | TATAAT |
| minus10_12G | GATAAT |
| minus10_12A | AATAAT |
| minus10_7A | TATAAA |

**Supplementary Table 5) -10 element sequences used in this work.** These -10 sequences for RNAP recognition were previously reported and result in a wide range of binding affinities.

| UP element name | Sequence (5’>3’) |
|---|---|
| up_326x | GGAAAATTTTTTTCAAAAGTA |
| up_136x | GAAAATATATTTTTCAAAAGTA |
| up_69x | AGAAAATTATTTTAAATTTCCT |
| no_up | AGCTCATTCATTAGGCACCCCA |

**Supplementary Table 6) UP element sequences.** UP elements used in this work to generate function promoters lacking -35 elements.

| Primer | Sequence (5’>3’) |
|---|---|
| GU 59 | CATGTTGTCCACTCCAATCGGTGATGGTCCTG |
| GU 60 | GTAATAGCTAAATCCCACCCGATGCCTGCAGG |
| GU 65 | CAAGCAGAAGACGGCATACGAGAT CGAATG CATGTTGTCCACTCCAATCG |
| GU 66 | CAAGCAGAAGACGGCATACGAGAT CTATGC CATGTTGTCCACTCCAATCG |
| GU 67 | CAAGCAGAAGACGGCATACGAGAT GCTAGT CATGTTGTCCACTCCAATCG |
| GU 68 | CAAGCAGAAGACGGCATACGAGAT GTACTG CATGTTGTCCACTCCAATCG |
| GU 70 | AATGATACGGCGACCACCGAGATCTACACGTAATAGCTAAATCCCACCCGATGC |
| GU 79 | CGTGCATAGTGCCATGTTATCCCTGAAGTCGAG |
| GU 86 | CAAGCAGAAGACGGCATACGAGAT GCTAGT CGTGCATAGTGCCATGTTATC |
| GU 87 | CAAGCAGAAGACGGCATACGAGAT GTACTG CGTGCATAGTGCCATGTTATC |
| GU 89 | CATAGCCGAATAGCCTCTCCACC |
| GU 99 | GCGATTGGTCTCACTAGAGCTGTC |
| GU 100 | GGTCAGCCATGGTTATTTGTACAGTTC |
| GU 102 | AATGATACGGCGACCACCGAGATCTACAC |
| GU 132 | TGTCAGGCATATTATCCGCT |
| GU 133 | CGGTTTATGGGTGTTATCGC |
| GU 134 | TCGTATCCCTGCAGGNNNNNNNNNNNNNNNNNNNNGCATGTGAGACCCGGTTTAT GGGTGTTATCGC |
| GU 142 | GGTCCAGTGCCATGTTATCCCTGAAGT |

**Supplementary Table 7) Primers used in the study.** See methods for description of primer usage.

| Library name | Uninduced expression | Induced expression | Fold-change |
|---|---|---|---|
| Pcombo<br>(N = 1493) | Min: 0.284<br><br>Max: 75.7<br><br>Range: 267x | Min: 0.126<br><br>Max: 57.0<br><br>Range: 453x | Min: 0.222<br><br>Max: 8.97<br><br>Range: 40.4x |
| Pspacer<br>(N = 3769) | Min: 0.0838<br><br>Max: 193<br><br>Range: 2300x | Min: 0.0567<br><br>Max: 183<br><br>Range: 3230x | Min: 0.132<br><br>Max: 15.6<br><br>Range: 118x |
| Pmultiple<br>(N = 1638) | Min: 0.0888<br><br>Max: 85.6<br><br>Range: 963x | Min: 0.159<br><br>Max: 74.3<br><br>Range: 467x | Min: 0.174<br><br>Max: 16.6<br><br>Range: 95.2x |
| Psteric<br>(N = 1369) | Min: 0.164<br><br>Max: 33.3<br><br>Range: 202x | Min: 0.135<br><br>Max: 23.3<br><br>Range: 173x | Min: 0.217<br><br>Max: 11.8<br><br>Range: 54.3x |

**Supplementary Table 8) Library range statistics.** Reported statistics amongst all promoters characterized in each library.
